# Structural evolution of Fcα/μR-mediated antibody recognition

**DOI:** 10.64898/2026.11.06.764962

**Authors:** Madhusree Seal, Walter N. Harrington, Ekaterina M. Flaugher, Richard J. Webby, Beth M. Stadtmueller

## Abstract

The Fc alpha-mu receptor (Fcα/μR) is an ancient Fc-receptor sharing common ancestry with the polymeric Ig receptor (pIgR) and binding polymeric IgM and IgA ligands. In several vertebrate lineages, including birds, Fcα/μR may serve as the sole IgM and IgA Fc receptor; yet its evolution and ligand-recognition mechanisms across vertebrates remain poorly understood. Here, we identify *FCAMR* expression in duck (*Anas platyrhynchos*, *ap*) lymphoid tissues and demonstrate that recombinantly expressed *ap*Fcα/μR ectodomain binds *ap*IgM and *ap*IgA ligands. *ap*Fcα/μR affinity was ∼10-fold higher for *ap*IgM versus *ap*IgA and ∼10-fold higher than mouse (*mm*) Fcα/μR. Cryo-EM structures of *ap*Fcα/μR in complex with pentameric *ap*IgM (3.27 Å) and tetrameric *ap*IgA (3.57 Å) showed three Ig-like domains (D1–D3) bound asymmetrically with 1:1 stoichiometry. Comparisons to avian pIgR, which contains four Ig-like domains (D1-D4), revealed overlapping contacts between *ap*Fcα/μR-D1 and pIgR-D1. *ap*Fcα/μR D2–D3 engaged sites similar to pIgR D3–D4, yet formed distinct contact networks. Parallel analysis of a *mm*Fcα/μR—*mm*IgM structure (4.13 Å) revealed *mm*Fcα/μR’s single Ig-domain bound analogous to *ap*Fcα/μR-D1 and to mammalian pIgR-D1, whereas downstream ectodomain regions were not observed. Collectively, the data elucidate PIGR-family mechanisms across vertebrates, providing a foundation for delineating common and divergent functions.

## INTRODUCTION

Immunoglobulins (Igs), or antibodies, play a critical role in immunity by combining hypervariable antigen binding fragments (Fabs) with one constant fragment crystallizable (Fc). All jawed vertebrates express multiple antibody isotypes, allowing them to combine Fabs with different Fcs to direct antibodies to specific compartments and/or activate downstream effector functions. These immune regulatory processes rely on a diverse family of Fc receptors (FcRs), each of which are expressed on specific cell types and selectively recognize different isotypes. While a subset of FcRs induce antibody transcytosis, others induce effector functions such as antibody-mediated phagocytosis, antigen processing and presentation, antibody-dependent cell-mediated cytotoxicity, and cytokine release, all critical to combating disease.

From an evolutionary standpoint, the oldest known FcRs belong to the polymeric immunoglobulin receptor (*PIGR*) family, which includes polymeric immunoglobulin receptor (pIgR; encoded by *PIGR*), Fc alpha and mu receptor (Fcα/μR; encoded by *FCAMR*), and Fc mu receptor (FcμR; encoded by *FCMR*)^1^. Although pIgR has been characterized across bony fish, amphibians, birds, reptiles, and mammals, its presence in bony fish may have arisen through convergent evolution^2,3^. Amphibians exhibit the first clear evidence of Fcα/μR and pIgR, which are cis duplicates arising from a common ancestral gene^1^, whereas FcμR emerged in reptiles^4^. The *PIGR* family synteny is conserved from amphibians to mammals (tetrapods) where FCAMR is found adjacent and upstream of PIGR^1,5^. However, Fcα/μR structures and functions remain poorly characterized across species, especially in birds.

All members of the PIGR family are type 1 transmembrane proteins that utilize one or more extracellular Ig-like domains to bind ligands. The Ig domain is a common ∼100-residue structural fold packed into two β sheets that form a β-sandwich stabilized by a disulfide bond. Ig domains are classified based on the number and location of β-strands. Most *PIGR* family receptors contain one or more Ig-Variable (V) type domains^3^, which possess β strands A, B, E and D on one sheet and A′, G, F, C, C’ and C” strands on the other^6^. Their ligands are polymeric (p) Igs, which contain two to six antibody monomers and up to one joining chain (JC); pIgs span multiple isotypes including pIgM and pIgA, and in amphibians, pIgX, an IgA ancestral analog^7^. JC is necessary for pIgR binding^8–12^ and reportedly contributes significantly to Fcα/μR binding^1,13^. Both receptors engage the IgM Fc (Fcμ; domains Cμ2–Cμ4 in mammals and birds) and/or the IgA Fc (Fcα; domains Cα2–Cα3 in mammals and Cα2–Cα4 in birds).

Functionally, the pIgR is expressed on epithelial cells where it binds and transcytoses pIgM and pIgA to mucosal secretions. During transcytosis, the ectodomain, called secretory component (SC), is cleaved and remains covalently attached to the pIg when released, forming secretory (S) antibodies. In contrast, Fcα/μR is expressed on both non-hematopoietic and hematopoietic immune cells and in mammals mediates pIgM and pIgA antigen immune complex endocytosis^13–15^. Among mammals Fcα/μR tissue distribution varies, with mouse Fcα/μR being expressed on a subset of B cells and macrophages in spleen^13–18^, kidney^14–16^, intestine^14^, and lymph nodes^16,19^ and human Fcα/μR also being expressed in tonsils^20^ but not in the spleen^15,20^. Within these tissues, Fcα/μR appears to be enriched in germinal centers, especially on the surface of follicular dendritic cells and marginal zone B cells residing in germinal centers^17,18,20–22^.

Structural studies since 2020 have revealed numerous pIg structures from bony fish, amphibians, birds and mammals, including their complexes with SC and FcμR^23^; however, Fcα/μR structures and complexes with ligands have remained unreported, hindering our understanding of this ancient receptor and its relationship to *PIGR* family members. Here we examine the structural evolution of Fcα/μR, first identifying *FCAMR* gene expression in birds and subsequently determining avian and mammalian Fcα/μR—IgM complex structures as well as the avian Fcα/μR—IgA complex structure. Collectively, these data elucidate Fcα/μR structural similarities and differences and interacting mechanisms across vertebrates, while also facilitating comparisons among PIGR family receptors relevant to understanding immune system evolution and FcR functions.

## RESULTS

### Conserved genomic architecture and tissue expression of avian *FCAMR*

We conducted a comparative analysis of the human and mallard duck (*Anas platyrhynchos*; *ap*) genomic assemblies, which revealed that the *FCAMR-PIGR* locus is evolutionarily conserved as a syntenic block across human chromosome 1 and mallard chromosome 27. Consistent with previous reports^1^, we did not find evidence for the *FCMR* gene within the *PIGR* family locus in birds. Adjacent to *PIGR*, we identified a putative *ap FCAMR* ortholog (Fig. 1a). While the mammalian Fcα/µR ectodomain has one immunoglobulin (Ig)-like domain followed by a predicted unstructured region, domain prediction identified three Ig-like domains within the *ap* Fcα/µR ectodomain (Fig. 1b), similar to *Xenopus* Fcα/µR^1^.

**Fig. 1.**
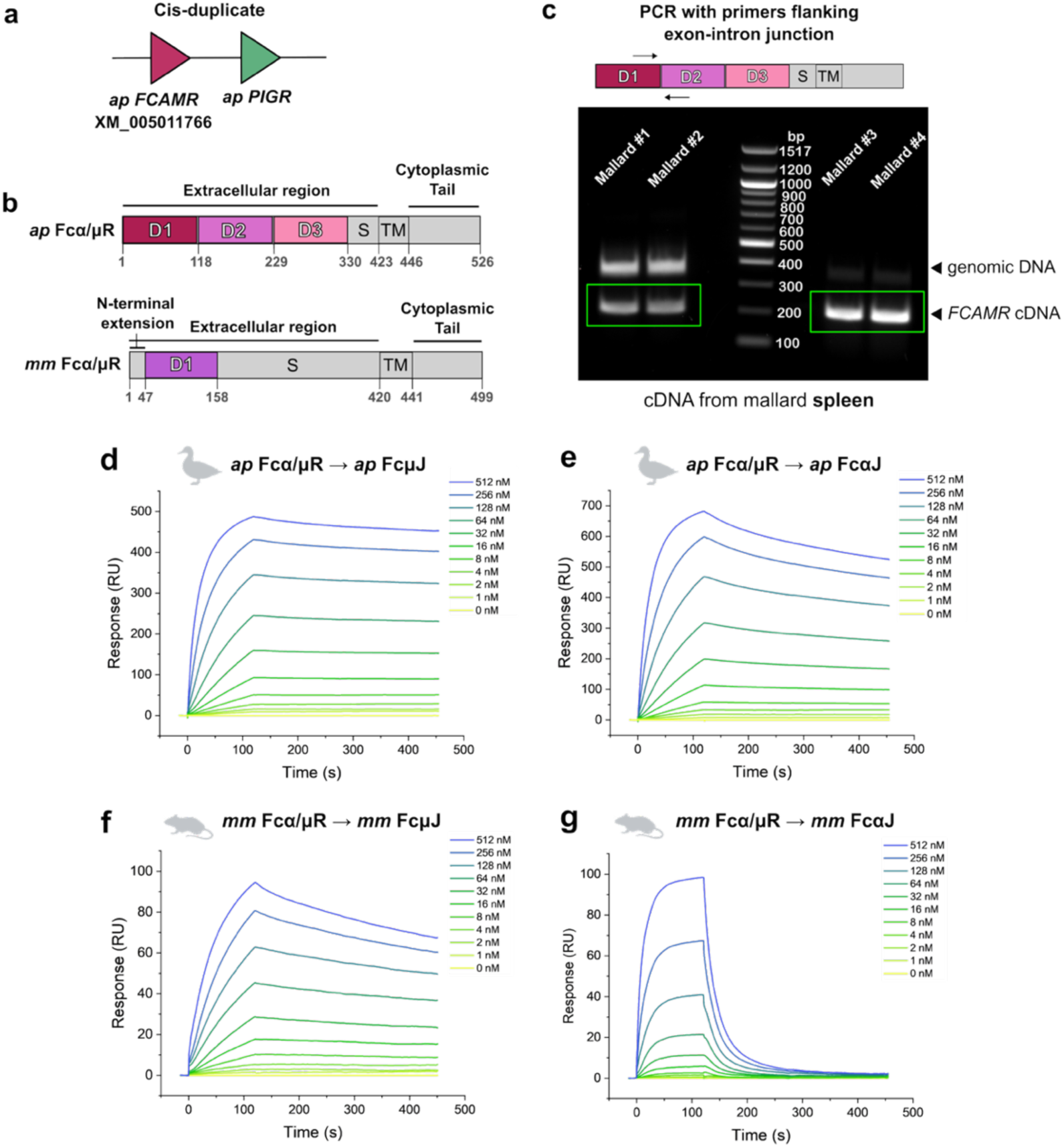
Genomic organization, domain architecture, and binding kinetics of mallard and mouse Fcα/μR. **(a)** Schematic representation of the PIGR family locus on the mallard genome (GCF_047663525.1) showing the tandem arrangement of the mallard (*Anas platyrhynchos*, *ap*) *FCAMR* gene (XM_005011766) and *ap PIGR* on chromosome 27. **(b)** Schematic comparison of the domain organization of *ap* (top) and mouse (*Mus musculus*, *mm*; bottom) Fcα/μR. *ap* Fcα/µR’s three Ig-like domains (D1-D3), stalk region (S), transmembrane region (TM), and cytoplasmic tail are indicated. Similarly, *mm* Fcα/µR’s unstructured N-terminal extension, Ig-like domain (D1), S region, TM region and cytoplasmic tail are indicated. Numbering begins at the residue following the signal peptide. The extracellular region was recombinantly expressed for our studies. **(c)** Schematic (top) showing the primer design flanking the *ap FCAMR* D1–D2 exon-intron junction across four biological replicates (Mallard #1-4). Black arrowheads indicate the genomic DNA product (406 bp) and the sequence-verified *ap FCAMR* cDNA product (229 bp). Green boxes highlight the expression of *ap FCAMR* across four biological replicates. A molecular weight ladder is shown in the center. (**d-g**) SPR sensorgrams showing response units (RU) for a two-fold dilution series of: (**d**) *ap* Fcα/µR binding to immobilized *ap* FcµJ, (**e**) *ap* Fcα/µR binding to immobilized *ap* FcαJ, (**f**) *mm* Fcα/µR binding to immobilized *mm* FcµJ, and (**g**) *mm* Fcα/µR binding to immobilized *mm* FcαJ. In **d-g**, each analyte response is colored according to the key on the right.

To confirm *FCAMR* tissue expression in mallard ducks, we isolated RNA from spleen, small intestine, bursa of fabricus, cloaca, and esophagus, and performed RT-PCR with intron spanning primers. Initial PCR amplification of spleen cDNA revealed a 229-base pair (bp) *FCAMR* transcript amplicon alongside a 406-bp product (Fig. 1c). DNA sequencing confirmed that the 229-bp and 406-bp amplicons corresponded to *FCAMR* mRNA and genomic DNA, respectively (Supplementary Table 1). Equivalent experiments also detected *FCAMR* expression in the intestine, bursa and cloaca, but not in the esophagus (Supplementary Fig. 1a-d).

### Fcα/µR ectodomain binds cognate ligands

To confirm that the putative *ap FCAMR* gene forms a functional protein that binds IgA and IgM, we designed a recombinant expression construct encoding the Fcα/µR ectodomain (hereafter *ap* Fcα/µR), which we produced and purified along with *ap* pIgM and pIgA variants lacking Fabs (hereafter *ap* FcµJ and *ap* FcαJ, respectively). For comparison we also produced mouse (*mm*) Fcα/µR ectodomain (hereafter *mm* Fcα/µR) and *mm* pIgM and pIgA variants lacking Fabs (hereafter *mm* FcµJ and *mm* FcαJ). Subsequently, we assayed binding between Fcα/µR analytes and cognate FcµJ and FcαJ ligands using Surface Plasmon Resonance (SPR) binding assays. *ap* Fcα/µR exhibited concentration dependent binding to both *ap* FcµJ (Fig. 1d) and *ap* FcαJ (Fig. 1e) with moderate association and slow dissociation. Consistent with prior SPR experiments measuring SC binding to pIgM and pIgA ligands^24–26^, *ap* Fcα/μR responses did not fit single-state kinetic models. Therefore, we analyzed the responses using EVILFIT software, which accommodates deviations from a standard 1:1 Langmuir interaction model by resolving continuous distributions of affinity and rate constants to determine an approximate K_D_^27–30^. EVILFIT-derived values suggested an approximate 10-fold difference in affinity between pIgM and pIgA binding with *ap* Fcα/µR binding to FcµJ with a K_D_ ∼10^−9^ M, and to FcαJ with a K_D_ of ∼10^−8^ M (Supplementary Fig. 1e). Equivalent SPR experiments testing *mm* Fcα/µR binding to *mm* FcµJ and *mm* FcαJ also revealed robust concentration dependent binding, albeit with a more rapid dissociation compared to *ap* Fcα/µR binding to *ap* FcµJ and *ap* FcαJ (Figs. 1f, 1g). EVILFIT derived values indicated *mm* Fcα/µR binding to *mm* FcµJ and *mm* FcαJ with a K_D_ of ∼10^−8^ M and ∼10^−7^ M, respectively (Supplementary Fig. 1f). Together, results confirm putative *ap* Fcα/µR binding to both *ap* pIgM and pIgA, which is consistent with mammalian counterparts. However, data also indicate differences in *ap* Fcα/µR binding to *ap* pIgM versus *ap* pIgA and potential species-specific differences between mallards and mammals.

### Structures of avian and mammalian Fcα/µR complexes

To investigate differences between avian and mammalian Fcα/µR, we determined structures of *ap* Fcα/µR*–ap* FcµJ (Fig. 2a, b, Supplementary Fig. 2), *ap* Fcα/µR—*ap* FcαJ (Fig. 2c, d, Supplementary Fig. 3, 4), and *mm* Fcα/µR*—mm* FcµJ (Fig. 2e, f, Supplementary Fig. 5) complexes to resolutions of 3.27 Å, 3.57 Å, and 4.13 Å, respectively.

**Fig. 2.**
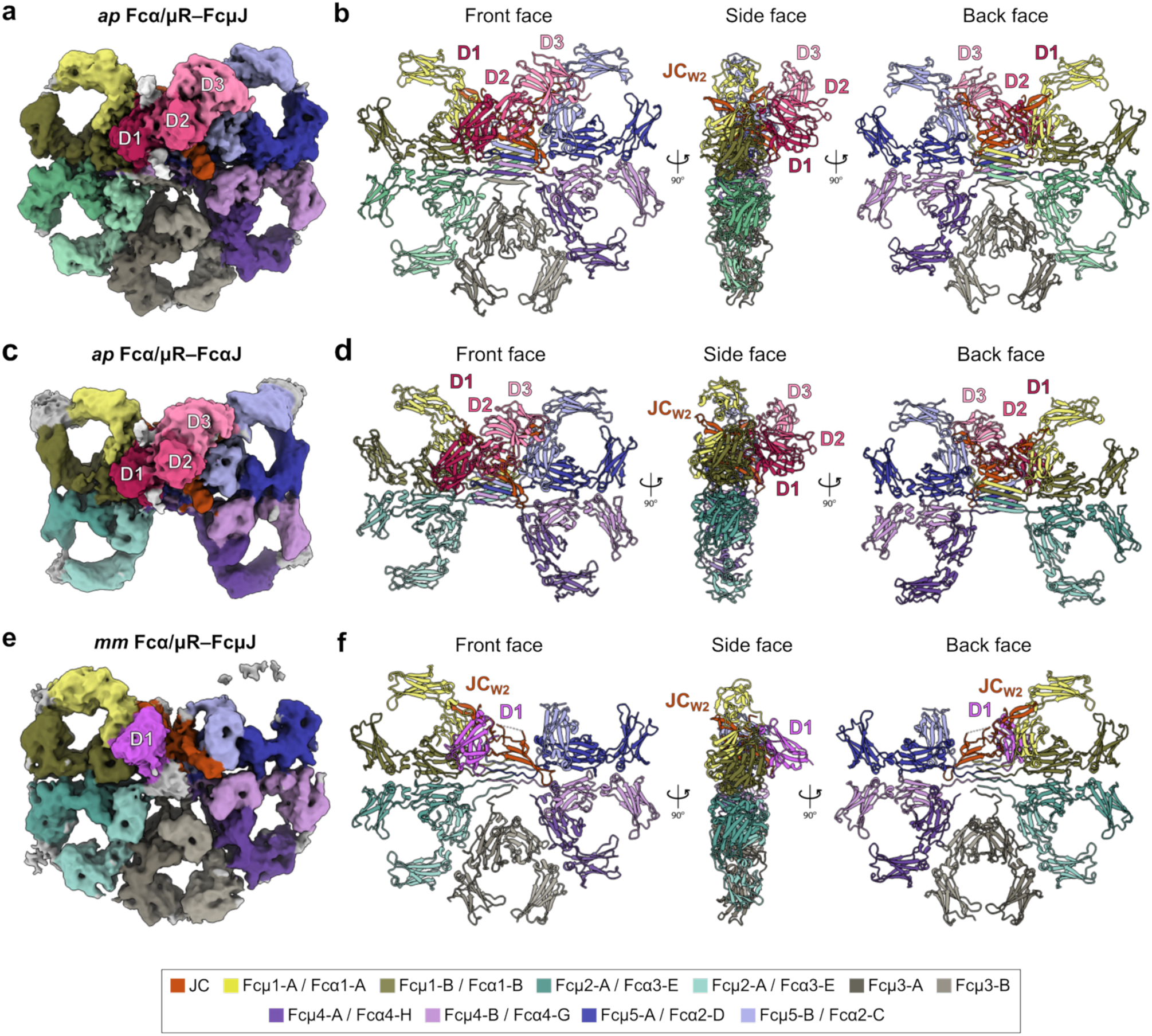
Structure of *ap* Fcα/µR–*ap* FcµJ, *ap* Fcα/µR–*ap* FcαJ and *mm* Fcα/µR–*mm* FcµJ. **(a, b)** Structure of *ap* Fcα/µR–*ap* FcµJ shown as (**a**) cryo-EM density map and (**b**) cartoon representation in three orientations (front, side and back faces). Density and cartoon models are colored by subunit according to the key below, with *ap* Fcα/µR domains D1, D2 and D3 colored in distinct shades of pink. **(c, d)** Structure of *ap* Fcα/µR–*ap* FcαJ shown as (**c**) cryo-EM density map and (**d**) cartoon representation in three orientations (front, side and back faces). Density corresponding to each chain is colored uniquely according to the key below and *ap* Fcα/µR domains (D1-D3) are colored in different shades of pink as in (a, b). **(e, f)** Structure of *mm* Fcα/µR–*mm* FcµJ shown as (**e**) cryo-EM density map and (**f**) cartoon representation in three orientations (front, side and back faces). Density and cartoon models are colored by subunit according to the key below, with *mm* Fcα/µR colored in fuchsia.

In the *ap* Fcα/µR–FcµJ structure, local resolution was variable, ranging from 3 Å to 6 Å (Supplementary Fig.2b) with well-resolved density for most main chain and side atoms at the interface. The distal domains including IgM Cµ2, were poorly resolved, consistent with reported flexibility observed in other structure containing FcµJ^25^. The refined *ap* Fcα/µR–FcµJ structure revealed three Ig-like domains asymmetrically bound to FcµJ with 1:1 stoichiometry (Fig. 2b), forming a 1732 Å^2^ interface with FcμJ. The 92 residues following D3 (corresponding to the glycosylated, mucin-like stalk region) were poorly resolved and not built. The FcµJ core was superimposable with previously reported structures, revealing five Fc subunits (Fcμ1 to Fcμ5, each comprising two HCs) and one JC arranged with distinct front and back faces (Fig. 2b)^25^. JC’s two β-hairpin wings (JC_W1_ and JC_W2_) were ordered in *ap* Fcα/µR—FcµJ, as observed in *ap* SFcµJ (PDB: 11DN).

Like *ap* Fcα/µR–FcµJ, the refined *ap* Fcα/µR—FcαJ structure revealed three Ig-like domains asymmetrically bound to FcαJ in 1:1 stoichiometry (Fig. 2d) with a 1303 Å^2^ interface. Local resolution was variable, ranging from 3 Å to 8 Å (Supplementary Fig. 4a) with well-resolved density for most main chain and side atoms in *ap* Fcα/µR D1 and at FcµJ interfaces. Distal domains, including the FcαJ Cα2 domain were poorly resolved, consistent with reported flexibility^24^. Despite well-ordered D1 density, D2 and D3 domains exhibited lower local resolution (5-8 Å) and evidence of variable occupancy compared to equivalent domains in *ap* Fcα/µR–FcµJ (Supplementary Fig. 4a). Accordingly, we refined main chain atom positions and built the side chains in geometrically reasonable positions. The FcαJ core was superimposable with previously reported structures^24^, revealing four Fc subunits (Fcα1 to Fcα4, each comprising two HCs) and one JC arranged with distinct front and back faces (Fig. 2d).

In the *mm* Fcα/μR–FcμJ structure, local resolution ranged from 4 Å to 9 Å. This limited overall side chain density, but the mainchain density and density for key interactions at the *mm* Fcα/μR–FcμJ interface were reliably resolved (Supplementary Fig. 5e-g). Accordingly, we refined the mainchain coordinates and placed the side chains in geometrically reasonable positions where visible density was poor. One copy of Cµ3, in chain Fcµ5-B (Fig. 2e), was poorly resolved as observed in prior mammalian IgM structures^31^, and therefore, not built. The refined *mm* Fcα/µR—*mm* FcµJ structure revealed one Ig-like domain asymmetrically bound to FcµJ, forming a 1324 Å^2^ interface. FcµJ was largely superimposable with human FcµJ found in published structures^31–33^. Unlike *ap* Fcα/µR–FcµJ, JC_W1_ was disordered in *mm* Fcα/µR—FcµJ, consistent with a subset of published structures containing mammalian FcµJ^31–33^.

The three structures share numerous commonalities and differences. For example, in all structures, the Fcα/µR D1 domain uses loops structurally equivalent to antibody complementarity determining regions (CDRs) 1-3 to form similar interfaces with JC and heavy chains (HCs). In contrast, Fcα/µR D2 and D3 are unique to *ap* Fcα/µR structures and use regions outside CDRs to form unique contacts with ligands. Accordingly, we first analyzed *ap* Fcα/μR D1 and *mm* Fcα/μR D1 interactions with FcμJ and then analyzed *ap* Fcα/μR D1 with FcαJ, comparing species and isoform-specific differences. Subsequently we compared Fcα/µR D2 and D3 interactions with FcμJ and FcαJ.

### Mallard and Mouse Fcα/µR-D1 interfaces with FcµJ

Structural comparison between *ap* Fcα/µR–FcµJ and *mm* Fcα/µR—FcµJ complexes revealed the Fcα/µR D1 domains using CDR-like loops (hereafter CDRs) 1-3 to contact JC, as well as Cμ4 domains in chains Fcμ1-B, Fcμ5-A and Fcμ5-B (Supplementary Fig. 6a). However, distinct from *ap* Fcα/µR, *mm* Fcα/µR D1 additionally binds to the Fcμ1A Cμ4 domain. In *ap* Fcα/µR, the CDR1 loop primarily contacts Fcµ1-B, JC loop residues between JC_W1_ and JC_W2_, and the JC C-terminus (Fig. 3b). Notable interactions between CDR1 and the JC C-terminal include putative hydrogen bonding between Lys29 and JC Asp138, Ser27 and JC Ala133 main chain atoms, and π-π stacking interactions between His30 and JC Tyr136. Additionally, CDR1 residues Asn28 and Asp31 contact JC Arg107 in the loop between JC_W1_ and JC_W2_ via hydrogen bonding and salt bridge interactions, respectively. Beyond these JC contacts, CDR1 Arg32 forms a salt bridge with Fcµ1-B Glu340. Residues immediately downstream of CDR1 were also observed forming interactions with Fcµ1-B as well as with Fcµ4-B, Fcµ5-A and Fcµ5-B Tp residues. For example, Arg38 in the D1 C β-stand contacts Fcµ5-A and Fcµ4-B Tp main chain (Supplementary Fig. 6a). Furthermore, Asn41, Gly42 and Val43 in CC’ loop form a tight van der Waals packing interface with Ala444 in Fcµ5-A, while Tyr45 and Thr46 mediate hydrophobic and hydrogen bonding interactions with residues Glu398 and His339, respectively, in Fcµ1-B.

**Fig. 3.**
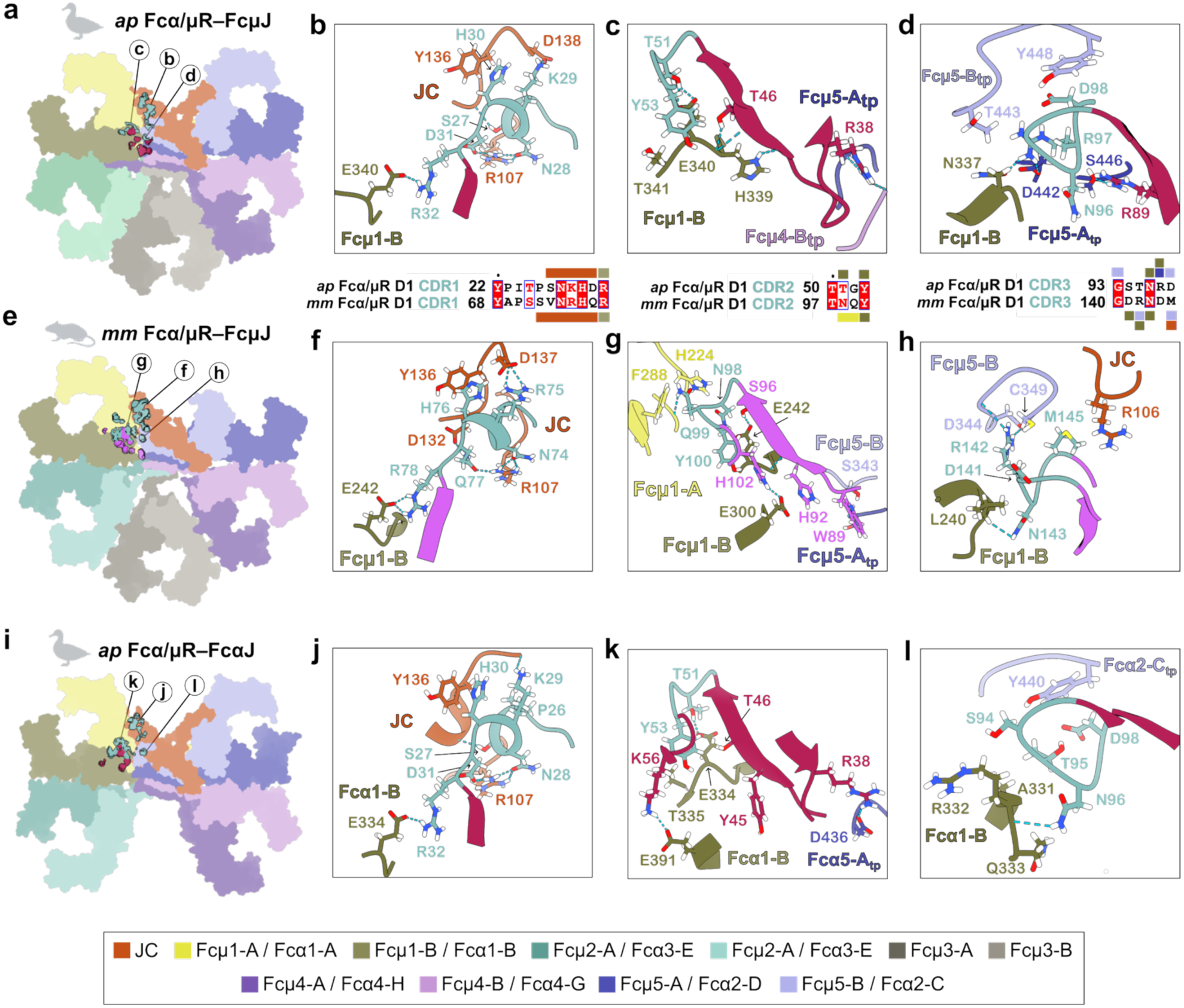
*ap* Fcα/µR D1 and *mm* Fcα/µR D1 interfaces with cognate FcµJ and FcαJ. **(a)** *ap* FcµJ shown as a flat surface with the *ap* Fcα/µR D1 contact footprint shown in cyan (CDR residues) and raspberry (residues outside CDRs). Circled letters denote interfaces magnified in **b-d**. **(b-d)** Top: Closeup views of *ap* Fcα/µR D1 (**b**) CDR1, (**c**) CDR2, and (**d**) CDR3 interfaces with *ap* FcµJ; contacts are shown as sticks, with putative hydrogen bonds/salt bridges indicated by dashed lines. Bottom: structure-based sequence alignments comparing *ap* Fcα/µR D1 CDRs with *mm* Fcα/µR D1 CDRs; contact residues and interacting chains are indicated with color-coded boxes. **(e)** *mm* FcµJ shown as a flat surface with the *mm* Fcα/µR D1 contact footprint shown in cyan (CDR residues) and fuchsia (residues outside CDRs). Circled letters denote interfaces magnified in **f–h**. (**f-h**) Closeup views of *mm* Fcα/µR D1 (**f**) CDR1, (**g**) CDR2, and (**h**) CDR3 interfaces with *mm* FcµJ; contacts are shown as sticks, with putative hydrogen bonds/salt bridges indicated by dashed lines. **(i)** *ap* FcαJ shown as a flat surface with the *ap* Fcα/µR D1 contact footprint shown in cyan (CDR residues) and raspberry (residues outside CDRs). Circled letters denote interfaces magnified in **j–l**. (**j-l**) Close-up views of *ap* Fcα/µR D1 (**j**) CDR1, (**k**) CDR2, and (**l**) CDR3 interfaces with *ap* FcαJ; contacts are shown as sticks, with putative hydrogen bonds/salt bridges indicated by dashed lines.

Adjacent to *ap* Fcα/µR CDR1, CDR2 and CDR3 formed extensive contacts with Fcµ1-B, Fcµ5-A, and Fcµ5-B. CDR2 Thr51 and Tyr53, like CDR1 Arg32, contacted Fcµ1-B Glu340, with Tyr53 also contacting Fcμ1-B Thr341 (Fig. 3c). Proximal to CDR2, Arg89 (F β-stand) formed a hydrogen bond with Fcµ5-A Ser446 (Supplementary Fig. 6a). In CDR3, Arg97 formed a salt bridge with Fcµ5-A tailpiece Asp442 and hydrogen bond with Fcµ1-B Asn337. Asp98 contributed to an anion-pi interaction with the aromatic ring of Fcµ5-B tailpiece Tyr448. Finally, Gly93 and Asn96 complemented the network by packing against Fcµ5-B Tyr448 and Fcµ1-B Ala338 back bone respectively (Fig. 3d).

Although the *mm* Fcα/µR–*mm* FcµJ structure was lower resolution and the position of some side chains were not reliably determined, structural alignments and structure-based sequence alignments revealed plausible similarities and differences between how *ap* Fcα/µR D1 and *mm* Fcα/µR D1 engage IgM. We found that *mm* Fcα/µR and *ap* Fcα/µR D1 CDR1 structures and ligand contacts were largely conserved and buried comparable surface area (1193 Å^2^ and 1299 Å^2^, respectively). In contrast, residues in the *ap* Fcα/µR D1 C β-stand and CC’ loop displayed more ligand contacts, including a Arg38 hydrogen bond with Fcμ5A mainchain backbone, than *mm* Fcα/µR (Fig. 3e, f, Supplementary Fig. 6a).

*mm* Fcα/µR D1 CDR2 buried a larger surface area (268 Å^2^) compared to *ap* Fcα/µR (95 Å^2^); this difference arises primarily from *mm* Fcα/µR CDR2 residues Asn98 and Gln99 contacting Fcµ1-A residues His224 and Phe288 (Fig. 3g). Variable CDR2 contacts between *ap* Fcα/µR and *mm* Fcα/µR may result from differences in Fc geometry between *mm* FcµJ and *ap* FcµJ, which appear to leave *mm* FcµJ Fcµ1-A more accessible. D1 CDR3 sequence is less conserved between *ap* Fcα/µR and *mm* Fcα/µR but buried comparable interface area (290 Å^2^ and 328 Å^2^, respectively) on FcµJ. Compared to *ap* Fcα/µR D1 CDR3, *mm* Fcα/µR CDR3 Arg142 formed a salt bridge with Fcμ5-B Asp344 alongside hydrogen bonds with Fcµ1-B and Fcµ5-B backbone residues. Additionally, Asp141 and Asn143 mediated packing interactions with Fcµ1-B, while Met145 contacts Fcµ5-B Thr348 and JC Arg106 (Fig. 3h). Structurally, *ap* Fcα/µR D1 buries a larger interface area with Fcµ5-A (250 Å²) than *mm* Fcα/µR (64 Å²). Conversely, *ap* Fcα/µR D1 buries less interface area with Fcµ1-A and JC (13 Å² and 309 Å², respectively) than *mm* Fcα/µR D1 (203 Å² and 420 Å², respectively).

### Comparison of ap Fcα/µR D1 interaction with IgA vs IgM

Structural comparisons between *ap* Fcα/µR—FcαJ and *ap* Fcα/µR–FcµJ complexes revealed a largely conserved mode of D1 binding that relied on CDR-like loops and formed 994 Å^2^ and 1193 Å^2^ interfaces, respectively. *ap* Fcα/µR D1 contacts with FcαJ were mostly conserved with those of FcµJ (Fig. 3i-l); however, variations in ap Fcα/µR D1 contacts with Fcα2-D or the equivalent FcµJ chain, Fcμ5-A, were observed (Supplementary Fig. 6b). In *ap* Fcα/µR—FcαJ, CC’ loop residues Asn41, Gly42 and Val43 bury only 18 Å^2^. In contrast, *ap* Fcα/µR—FcµJ residues Asn41, Gly42, and Val43 form extensive Van der Waals contacts with Fcµ5-A and bury 78 Å^2^. Additionally, in *ap* Fcα/µR— FcαJ, *ap* Fcα/µR CDR3 does not contact Fcα2-D, whereas in *ap* Fcα/µR–FcµJ, CDR3 was positioned within hydrogen bonding distance of Fcµ5-A Asp442. Collectively these local differences cause *ap* Fcα/µR D1 to bury a significantly smaller surface area with Fcα2-D (80 Å^2^), than with Fcµ5-A (223 Å^2^).

### *ap* Fcα/µR-D2–D3 and interactions with FcµJ and FcαJ

While *ap* Fcα/µR D1 formed the largest buried surface area on both *ap* FcµJ and *ap* FcαJ, Fcα/µR D2 and D3 also mediated significant interfaces. In both structures, D2 and D3 were oriented differently than D1, with CDRs pointing away from FcµJ, allowing residues outside the CDR loops (i.e. D2 loops between β-strands A’-B, C-C’, C’-D and E-F of D2 and the linker preceding the D3 A’ β-strand) to mediate interfaces with FcµJ and FcαJ involving equivalent JC and HC regions (Supplementary Fig. 6c). Despite related modes of binding, differences were apparent. For example, while Fcα/μR D2, the D2–D3 linker, and D3 buried 125 Å^2^, 164 Å^2^ and 150 Å^2^, respectively, on *ap* FcµJ, these regions buried 48 Å^2^, 68 Å^2^ and 148 Å^2^, respectively, on *ap* FcαJ. Differences in buried surface area were associated with RMSDs of 4.898 and 6.325 respectively, when comparing the D2 and D3 positions in FcαJ relative to positions in FcµJ (Supplementary Fig. 6d), collectively indicating moderate differences in how D2–D3 engage IgM and IgA ligands.

Analysis of *ap* Fcα/µR–FcµJ, which exhibited a higher resolution interface, revealed D2, the D2–D3 linker, and D3 contacting the *ap* FcµJ JC N-terminus (JC_N-ter_), JC_W1_ and Fcµ5-B Cµ4 domain (Fig. 4a-c). Charged residues in the D2 C’D and A’B loops and D2–D3 linker dominated interactions with the JC_N-ter_ and JC_W1_, with D2–D3 linker residue Arg225 forming several key interactions. Hydrophobic and charged residues in D3, including Phe229 and Lys324, engaged JC_W1_ and Fcµ5-B (Fig. 4c). In contrast, *ap* Fcα/µR—FcαJ did not reveal direct contacts between Fcα/µR D2 and FcαJ (Fig. 4d); however, D2 approached the JC_N-ter_ where contacts might be apparent at higher resolution (Supplementary Fig. 6e). The *ap* Fcα/µR D2–D3 linker and D3 interfaces with FcαJ were well resolved (Supplementary Fig. 6e), and similar to *ap* Fcα/µR—FcµJ, revealed charged and hydrophobic residues contacting JC_W1_ and Fcα2-C (Fig 4d); however, the interface was notably smaller and was characterized by a more minor role for *ap* Fcα/µR Arg225. Differences in interfacing residues are in part explained by differences in HC sequence; indeed, while D1 residues contact residues that are largely conserved between the µ and α HCs, D2–D3 contact HC residues with lower identity and similarity (Supplementary Fig. 7a).

**Fig. 4.**
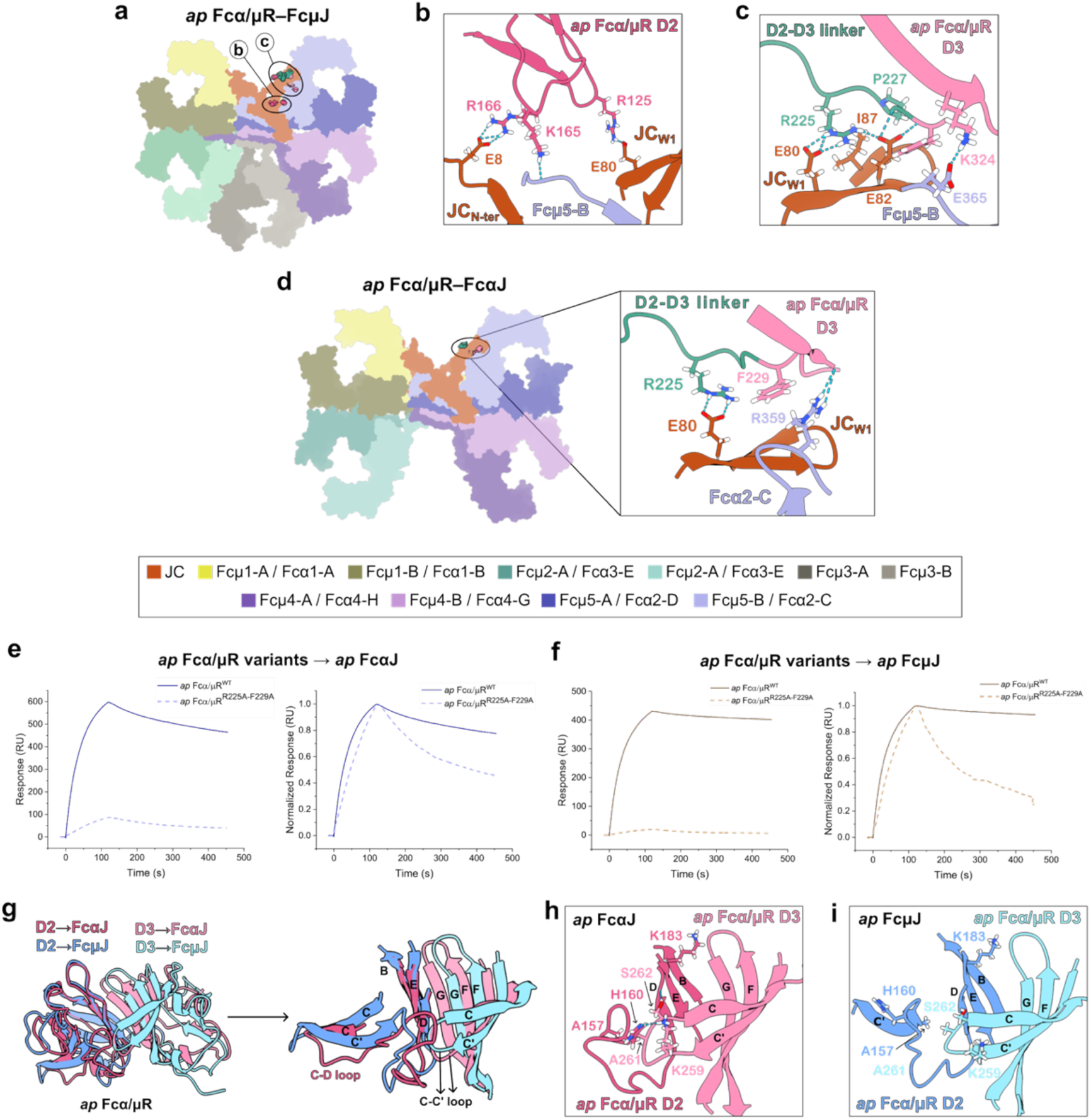
*ap* Fcα/µR D2–D3 interfaces with *ap* FcµJ and *ap* FcαJ. **(a)** *ap* FcµJ shown as a flat surface with the *ap* Fcα/µR D2 (dark pink), D2–D3 linker (cyan) and D3 (light pink) contact footprint shown. Circled regions denote interfaces magnified in **b** and **c**. **(b-c)** Detailed view of (**b**) *ap* Fcα/µR D2 interactions with the JC N-terminus (JC_N-ter_), JC_W1_ and Fcµ5-A, and (**c**) *ap* Fcα/µR D2–D3 linker and D3 interactions with the JC_W1_ and Fcµ5-A. Interacting residues are shown as sticks, with putative hydrogen bonds and salt bridges indicated by dashed lines. **(d)** *ap* FcαJ shown as a flat surface with the *ap* Fcα/µR D2–D3 linker (cyan) and D3 (light pink) contact footprint shown. The circled region is magnified (right) detailing the D2–D3 linker and D3 interactions with JC_W1_ and Fcα2-C. Putative hydrogen bonds and salt bridges are indicated by dashed lines. **(e, f)** SPR sensorgrams showing response units (RU, left) and normalized response (right) for 256 nM *ap* Fcα/μR variant analytes binding to (**e**) *ap* FcαJ or (**f**) *ap* FcµJ ligands. Solid lines indicate *ap* Fcα/μR while dashed lines indicate *ap* Fcα/µR^R225A-F229A^. Results are consistent with three or more replicate experiments. Sensorgrams for the full analyte concentration series and fitted data analysis of the *ap* Fcα/µR^R225A-F229A^ are provided in Supplementary Fig. 7b, d. **(g)** Structural superposition of *ap* Fcα/μR D2–D3 domains when bound to FcαJ (pink) versus FcµJ (blue), highlighting conformational shifts in the C–D and C–C′ loops. **(h, i)** Closeup view comparing the specific binding interfaces of *ap* Fcα/μR D2 and D3 with (h) *ap* FcαJ and (i) *ap* FcµJ. Key interacting residues are labeled and depicted as sticks.

To investigate how the *ap* Fcα/µR D2–D3 linker and D3 impact IgA and IgM binding, we designed and expressed a Fcα/µR variant, in which Arg225 and Phe229 were mutated to alanine (*ap* Fcα/µR^R225A-F229A^), and tested binding to *ap* FcαJ and *ap* FcµJ using SPR. Comparing concentration-matched responses and concentration-matched normalized responses, Fcα/µR^R225A-F229A^ exhibited decreased binding to both ligands compared to Fcα/µR, validating our structural observations and signifying a role for D2–D3 linker and D3 residues Arg 225 and Phe229 in ligand binding (Fig. 4e, f). Notably however, Fcα/µR^R225A-F229A^ binding to *ap* FcµJ exhibited a lower overall R_max_ and more rapid dissociation compared to Fcα/µR^R225A-F229A^ binding to FcαJ. EVILFIT derived values confirmed our qualitative observations, where *ap* Fcα/µR^R225A-F229A^ binding to FcαJ was associated with a K_D_ of ∼10^−7^ M (Supplementary Fig. 7b, c) and binding to FcµJ a K_D_ of ∼10^−5^ M (Supplementary Fig. 7d, e). Together data indicate that in *ap* Fcα/µR, Arg225 and/or Phe229 impacts interactions with IgM more than with IgA, a finding that may in part explain *ap* Fcα/µR’s higher affinity for IgM.

The observation that *ap* Fcα/µR formed fewer contacts with FcαJ compared to FcµJ suggested that other factors, such as *ap* Fcα/µR domain interfaces, might stabilize *ap* Fcα/µR interactions with FcαJ. Indeed, with respect to D1, D2 and D3 are orientated at 47° and 107° when bound to FcαJ compared to 38° and 87° when bound to FcμJ, leaving D3 closer to D2 when bound to FcαJ. While total buried surface area between D1 and D2 were similar in both structures, total BSA between D2 and D3 was higher (1522 Å^2^) in *ap* Fcα/µR—FcαJ compared to *ap* Fcα/µR–FcµJ (1166 Å^2^), suggesting that the positions of D2 and D3 are stabilized by additional inter domain contacts when bound to IgA. For example, in *ap* Fcα/µR—FcαJ, increased interface area arises in part from the D2 C-D loop shifting closer to the D3 C-C’ loop and enabling interactions that are absent in *ap* Fcα/µR—FcµJ (Fig. 4g-i). We hypothesize that together with additional inter-domain contacts, these interactions help to stabilize *ap* Fcα/µR interactions with IgA, where contacts with HC are limited.

### Comparisons between *ap* Fcα/µR and *ap* SC interactions with IgM and IgA

Given the evolutionary relationship between pIgR and Fcα/µR, including the common use of Ig domains to mediate interactions with IgA and IgM ligands, we compared Fcα/µR and SC containing structures. Of note, SC domain organization differs between mammals and birds, with mammals having five Ig domains (D1-D5) and birds having four domains (D1-D4) that are equivalent to mammalian D1, D3, D4 and D5, respectively^34^ (Fig. 5a, b). Structure and sequence-based alignment of individual *ap* Fcα/µR domains indicated that *ap* Fcα/µR D1, D2, D3 are equivalent to *ap* SC D1, D2, D3. Consistently, pairwise structural alignments between individual *ap* Fcα/µR and *ap* SC domains revealed RMSDs of 0.783 Å, 0.801 Å and 0.928 Å for D1, D2 and D3, respectively, indicating high structural similarity.

**Fig. 5.**
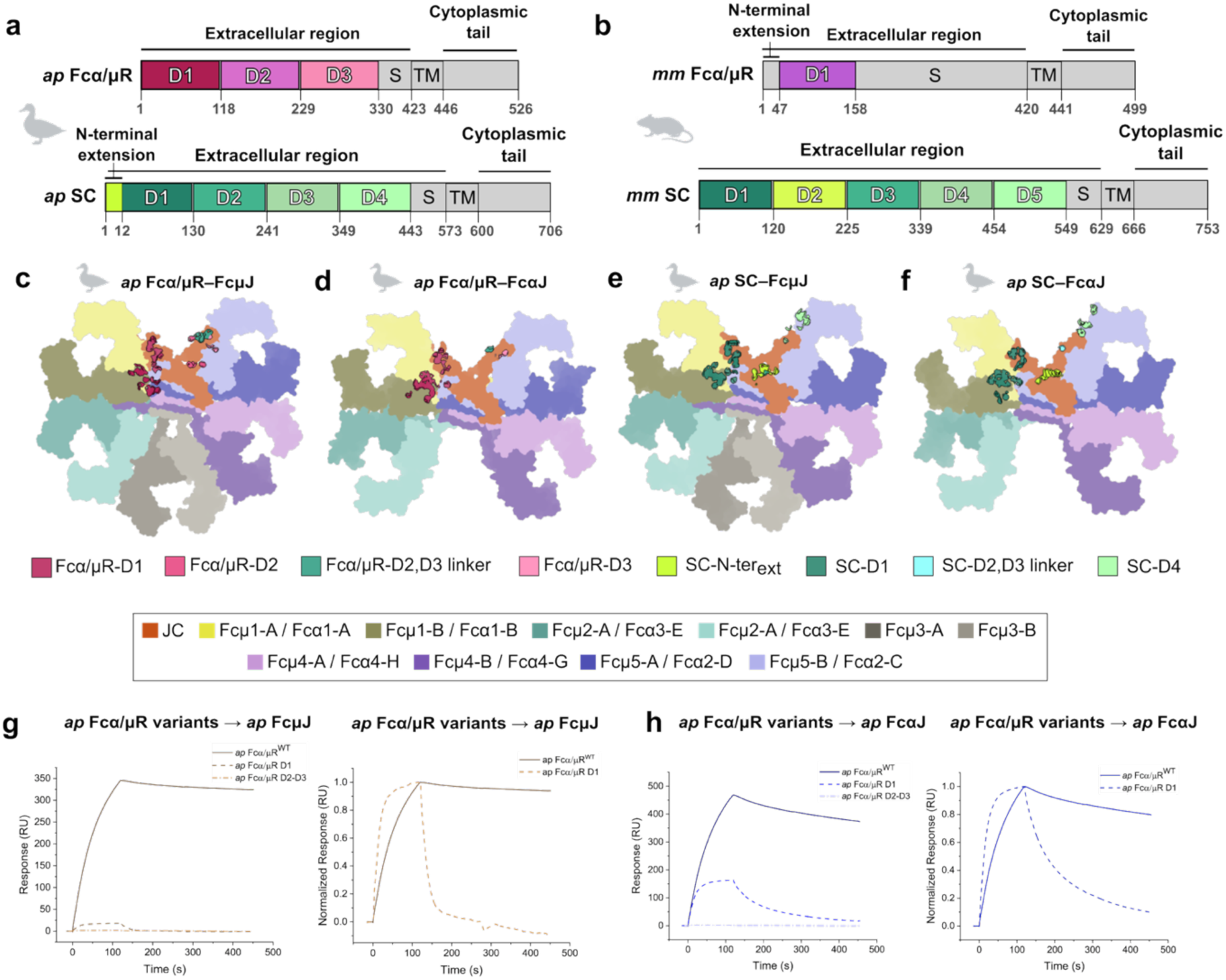
Structural comparison of *ap* Fcα/µR and *ap* SC interactions with IgM and IgA. **(a)** Schematic representation of *ap* Fcα/µR (top, shades of pink) and *ap* SC (bottom, shades of green) domain organization, indicating extracellular Ig-like domains (D1–D3 for Fcα/µR; N-terminal extension, D1–D4 for SC), S region, TM region, and cytoplasmic tail. **(b)** Schematic representation of mouse (Mus musculus, *mm*) Fcα/µR (top, fuchsia) and *mm* SC (bottom, shades of green) domain organization, indicating extracellular Ig-like domains (N-terminal extension, D1 for Fcα/µR; D1–D5 for SC), S region, TM region, and cytoplasmic tail. **(c-f)** Contact footprints showing *ap* FcµJ or *ap* FcαJ as flat surface representations with *ap* Fcα/µR or *ap* SC contacts indicated. Receptor domains, HCs and JC are colored according to the key below. **(g, h)** Surface Plasmon Resonance (SPR) sensorgrams showing response (left) and normalized response (right) for 256 nM *ap* Fcα/μR variant analytes binding to (**g**) *ap* FcµJ or (**h**) *ap* FcαJ ligands. Solid lines indicate *ap* Fcα/μR^WT^ while dashed lines indicate domain truncations (*ap* Fcα/µR D1 and *ap* Fcα/µR D2–D3). Sensorgrams for the full analyte concentration series and fitted data analysis of the *ap* Fcα/µR D1 are provided in Supplementary Fig. 10.

In both *ap* Fcα/µR and *ap* SC complexes with *ap* FcµJ and *ap* FcαJ, the interface is heavily dominated by D1, with *ap* Fcα/µR D1 burying ∼1193 Å² and *ap* SC D1 burying ∼1467 Å² on *ap* FcµJ. This difference is primarily mediated by an *ap* SC N-terminal extension (residues 1-11), without which the *ap* SC D1 buries only ∼955 Å² (Fig. 5c-f). The N-terminal extension allows *ap* SC to form additional contacts with the JC N-terminal residues (Fig. 5e, f). Conversely, *ap* Fcα/µR lacks an equivalent N-terminal extension and the first seven residues of JC are disordered in the *ap* Fcα/µR–FcµJ and Fcα/µR–FcαJ structures.

Both *ap* Fcα/µR D1 and *ap* SC D1 appear to share largely conserved modes of binding to *ap* FcµJ dominated by CDR1-3 contacts with JC, and Cµ4 domains of Fcµ1-B and Fcµ5-B (Supplementary Fig. 8a-i). However, some differences were apparent; for example, while *ap* Fcα/µR uses conserved CDR1 Arg32 to contact Fcµ1-B, the equivalent Arginine in *ap* SC CDR1 (Arg45) does not form contacts. A similar trend was observed while comparing *ap* Fcα/µR and *ap* SC binding to FcαJ, however unlike in *ap* SFcµJ, *ap* SC D1 CDR1 Arg45 contacts Fcα1-B (Supplementary Fig. 9a-i).

Observing conserved *ap* Fcα/µR D1 and *ap* SC D1 interfaces with ligands prompted us to test if *ap* Fcα/µR D1 was necessary and sufficient for binding to *ap* FcµJ and *ap* FcαJ ligands as has been shown for, mammalian SC D1 and Fcα/µR D1 ^26,35,36^. SPR experiments, which tested *ap* Fcα/µR, and D1- and D2–D3-only variants, revealed robust concentration dependent binding for *ap* Fcα/µR, *ap* Fcα/µR D1, but not *ap* Fcα/µR D2–D3 (a construct lacking D1). Comparing concentration-matched responses and concentration-matched normalized responses revealed that while *ap* Fcα/µR D1 is necessary for binding, D2–D3 slows dissociation thereby enhancing binding to both the ligands (Fig 5g, h).

Despite high structural conservation among individual domains and D1-ligand interactions, additional *ap* Fcα/µR and *ap* SC domains adopted distinct binding geometries on IgM and IgA. This was apparent when aligning *ap* FcαµR—FcµJ and *ap* SFcµJ structures on the JC, which yielded a low RMSD of 3.79 Å for D1, but higher RMSDs of 8.24 Å, and 12.59Å, for D2 and D3, respectively (Supplementary Fig. 10c). These differences were associated with differences in total buried surface area, with *ap* Fcα/µR and *ap* SC D2 burying 130 Å^2^ and 34 Å^2^, respectively, and *ap* Fcα/µR and *ap* SC D3 burying 220 Å^2^ and 76 Å^2^, respectively. Alignments comparing *ap* Fcα/µR–FcαJ and *ap* SFcαJ revealed similar trends (Supplementary Fig. 10d), where the positions of Fcα/µR and SC D2 and D3 were variable and Fcα/µR D2 and D3 buried a larger surface area.

Outside of D1, Fcα/µR and SC differences in buried surface area were associated with different contacts. For example, in *ap* Fcα/µR—FcµJ, *ap* Fcα/µR D2 residues Arg125, Arg166 and Lys165 contacted JC_W1_ Arg80, JC_N-ter_ Arg8 and Fcµ5-B main chain atoms, respectively, whereas in *ap* SFcµJ, *ap* SC D2 Glu204 forms only one probable hydrogen bond with JC_N-ter_ Arg10 (Supplementary Fig. 11b). Moreover, the noted role of the Fcα/µR D2–D3 linker, including Arg225 is more prominent than that of the SC D2–D3 linker, which forms limited interactions with JC_W1_ (Supplementary Fig. 11c). Similarly, *ap* Fcα/µR D3 Phe229 and Lys324 forms hydrophobic and hydrogen bond interactions with Fcµ5-B Glu365 whereas *ap* SC D3 has no contacts with FcµJ (Supplementary Fig. 11d). Inspecting contacts with FcαJ, revealed that Fcα/µR D2 and SC D2 both buried minimal surface area of 48 Å^2^ and 45 Å^2^, respectively, and that the Fcα/µR D2–D3 linker and SC D2–D3 linker contacts were similar to those observed in complexes with *ap* Fcµ (Supplementary Fig. 11e). Contact differences were more pronounced for D3; Fcα/µR D3 formed contacts with JC and Fcα2-C, whereas SC D3 formed no contacts with FcαJ (Supplementary Fig. 11f). While Fcα/µR lacks D4, SC D4 buried ∼382 Å^2^ interface area on FcµJ and FcαJ, and formed extensive contacts with JC_W1_ and Fcµ5-D/Fcα2-C including a covalent disulfide bond^24^. Together, while ap SC buries a larger total surface area compared to Fcα/µR, Fcα/µR relies more heavily on D2–D3 to augment D1 binding to IgM and IgA.

Given that *ap* Fcα/µR and *ap* SC inter-domain contacts, also appear to stabilize receptor-ligand interactions, we also compared interactions between D1 and D2. In *ap* SC, conserved residues form a hydrophobic interface, or clamp, between D1 and D2^24^, which is observed in mammalian SC structures as well^35,37^. Analysis of interdomain interfaces revealed that while *ap* Fcα/µR and *ap* SC D1–D2 interface surface area is similar (∼ 800-850 Å^2^), hydrophobic clamp residues, including ap SC D1 Pro14, Tyr22, and Leu24 and D2, Phe132 and Tyr133, as well as neighboring electrostatic interactions are not conserved in *ap* Fcα/µR, potentially providing more flexibility for Fcα/µR D2 and D3 to adopt variable positions when binding IgM versus IgA (Supplementary Fig. 11g).

### Mammalian Fcα/µR and pIgR ligand interactions

We also compared *mm* Fcα/µR D1 and mammalian SC D1 interactions with IgM. *mm* Fcα/µR and human (*hs*) SC D1 (reported in human SIgM structure, PDB: 7K0C) have an RMSD of 0.726, indicating high structural similarity. Although the structure of *mm* SC bound to *mm* IgM has not been reported and *mm* pIgR interactions with IgM are reportedly of lower affinity^2,38^, the high sequence similarity shared between *mm* and *hs* SC (77.2%) and IgM HC Cu4 (99.2%) make general comparisons reasonable. Aligning *mm* Fcα/µR–FcµJ and *hs* SFcµJ structures on the JC revealed an RMSD of 3.313 Å, similar to what we observed between *ap* Fcα/µR–FcµJ and *ap* SFcµJ. Additionally, structure-based sequence alignments indicate a highly conserved mode of binding between *mm* Fcα/µR D1, *ap* Fcα/µR D1, *mm* SC D1 and *ap* SC D1 (Supplementary Fig. 12a). Thus, the Fcα/µR and SC D1 domains retain largely conserved structure and ligand binding interactions whereas subsequent domains (e.g. ap Fcα/µR D2–D3, SC D2-D5, and mm Fcα/µR unstructured stalk region) have evolved unique attributes.

## DISCUSSION

Here we identify and characterize an avian Fcα/μR and report structures of *ap* Fcα/μR D1–D3 and *mm* Fcα/μR D1, which detail interactions with ligand and shed light on the evolution and function of PIGR-family FcRs across multiple species. Although the avian *FCAMR* gene was previously annotated in chickens (*Gallus gallus*)^1,39^, whether birds express a functional Fcα/μR capable of binding both IgM and IgA remained unverified. We confirmed *FCAMR* mRNA expression in mallard duck spleen, intestine, bursa of fabricus, and cloaca. Expression in duck lymphoid tissue is consistent with *FCAMR* expression data from amphibian *Xenopus laevis,* mice, and humans. While this points toward the existence of functional Fcα/μR in all birds, many avian genomes remain incompletely characterized, and lymphoid tissue organization is variable among bird species; thus, Fcα/μR prevalence and/or tissue distribution may vary. For example, most birds lack encapsulated lymph nodes whereas Anseriformes (waterfowl like ducks, geese, and swans) contain two pairs of organized lymph nodes localized in cervico-thoracic and lumbar regions^40,41^. While we did not characterize the Fcα/μR-expressing cells in ducks, in mammals Fcα/μR has been found on follicular dendritic cells, mesangial cells, a subset of B cells, and macrophages. Expression has been confirmed in mouse spleen, kidney, intestine, bone marrow, placenta, thymus^14^ and lymph nodes as well as in human tonsils, lymph nodes, and kidney^42^. Moreover, *Xenopus* Fcα/μR has been shown to be expressed in spleen^1^.

### Fcα/μR structures serve as models for functional receptors from birds and mammals

Mallard Fcα/μR utilizes three Ig-like domains (D1–D3) to interact with pentameric IgM and tetrameric IgA. In contrast, mouse *mm* Fcα/μR uses only a single Ig domain (D1) to bind IgM and IgA. Previous studies suggested that Fcα/μR domain organization and D1 sequence is conserved across mammalian species^14,20^. Our sequence analysis across representative avian species (Supplementary Fig. 12b) suggests that avian Fcα/μR domain organization and sequence is also largely conserved. Thus, *ap* Fcα/μR-FcµJ, *ap* Fcα/μR-FcαJ, and *mm* Fcα/μR-FcµJ containing structures are expected to serve as models for avian and mammalian Fcα/μR, respectively.

While our structures detail Fcα/μR – ligand interactions, it is important to consider that Fcα/μR is a type 1 transmembrane protein receptor expressed on the cell membrane surface. Structures of other type-I FcR ectodomains (e.g. pIgR, FcμR) are thought to serve as models for membrane bound receptors because the ectodomains typically comprise one or more structured domains (e.g. Ig-like domain) tethered to the TM via a flexible domain. Our work indicates that duck Fcα/μR ectodomain comprises a three-domain, D1–D3, ligand binding region and a disordered, 92-residue mucin-like stalk region containing 12 Serine and 25 Threonine residues; this connects to the TM, which is followed by an 81-residue intracellular region. In mammals the Fcα/μR ligand binding domains appear to comprise only D1 and a 262-residue mucin-like^43^ stalk region that is also predicted to be largely unstructured and to contain three Asparagine, 30 Serine and 31 Threonine residues that may serve as O-linked glycosylation sites.

While FcR’s structured ligand binding domains may be conformationally independent from linker or stalk regions, they may promote receptor clustering and/or intracellular, post-translational modifications that regulate signaling cascades. For example, the PIGR family receptor, FcμR (found only in mammals) comprises an ectodomain with one Ig-like domain and a 127-residue stalk region. Up to eight FcμR copies can bind a single IgM, which appears to promote interactions between stalk domains^23^. Although mammalian and avian Fcα/μR both contain mucin-like stalk regions, their ligand binding activity is most similar to pIgR, binding JC-containing antibodies with a 1:1 stoichiometry. Thus, one Fcα/μR stalk might interact with other stalks from Fcα/μR bound to adjacent copies of IgM or IgA, but interactions would be distinct from those observed when multiple FcμR bind a single IgM^23^. Differences in stalk length may also impact functions and/or the location of the ligand binding domain relative to the cell surface. Larger distances may help to accommodate large polymeric IgM/IgA-antigen immune complexes and/or may provide platforms for interactions between multiple stalk domains and/or binding partners. Indeed, prior reports suggest that mammalian Fcα/μR can form a heterodimer with an unknown protein^44^ and thus, the stalk region could act as a docking hub for the unidentified partner, but this hypothesis merits further investigation.

Regardless of stalk function(s) all Fc receptors within the *PIGR* family, including pIgR, Fcα/μR, and FcμR, have cytoplasmic tails that help regulate function. Whereas the pIgR cytoplasmic tail dictates transcytosis, the functions and mechanisms of Fcα/μR and FcμR remain less clear. In fact, all *PIGR* family receptor lack canonical intracellular signaling motifs such as immunoreceptor tyrosine-based activation motifs (ITAM), immunoreceptor tyrosine-based inhibition motifs (ITIM), and switch motifs, although tyrosine residues in FcμR cytoplasmic tail have been implicated in BCR signaling, anti-Fas induced apoptosis, and internalization^45^. The mouse Fcα/μR CT is known to contain a dileucine motif which promotes receptor internalization whereas in human Fcα/μR an internalization motif has not been identified. Our analysis of the mallard Fcα/μR sequence uncovered a YXXΦ motif, which is known to induce clathrin-mediated endocytosis, like the di-leucine motif^46,47^ and thus we speculate that receptor internalization maybe a common Fcα/μR function among species. Mouse and human Fcα/μR can internalize antigen-bound IgM and IgA, and thus internalization may promote antigen uptake functions, yet our SPR data suggests that *mm* Fcα/μR and *ap* Fcα/μR can bind to IgM and IgA with high affinity in the absence of antigen. Further studies are needed to define variables that regulate Fcα/μR mechanisms.

### Avian and mammalian Fcα/μR divergence

Why avian and mammalian Fcα/μR evolved different domain organization is not entirely clear. Birds maintain a three-Ig-like domain scaffold (D1–D3) that stabilizes high-affinity antibody engagement and mammals appear to accomplish this with a single domain; however, our SPR data indicate that avian Fcα/μR binds both ligands with 10-fold higher affinity than mouse Fcα/μR, suggesting that higher affinity binding may be advantageous in birds. Yet in both species, there appear to be similar differences in affinity for IgM and IgA ligands. Previous studies reported mammalian Fcα/μR to bind to both IgM and IgA, but with a 10-fold higher affinity for IgM. Consistent with these observations, we found that that mallard Fcα/μR bound to IgM and IgA, with IgM having approx. 10-fold higher affinity. The divergence in ectodomain architecture between avian and mammalian Fcα/μR might reflect distinct immunological and/or biophysical demands across vertebrate classes. Lacking FcμR and FcαRI^1,3^, birds may rely on Fcα/μR as their primary, classical receptor for both IgM and IgA whereas mammals, possessing an expanded repertoire of isotype-specific Fc receptors, may rely on distributed functions and/or redundancy among receptors. Notably, among three antibody isotypes (IgM, IgA and IgY) produced by birds, mallards primarily produce an Fc truncated version of IgY (IgYΔFc) in response to antigens, which is not known to bind classical FcRs. Thus, we hypothesize that Fcα/μR in mallards, and perhaps other birds, play a significant role in inducing antibody-mediated FcR effector functions.

### PIGR Family

More broadly, our findings shed light on key structural and mechanistic differences among the PIGR family members, in particular how Fcα/μR occupies a distinct functional niche relative to the pIgR/SC and FcμR. Although both cis-duplicates Fcα/μR and pIgR employ a closely related mode of ligand binding, there are notable differences in their structures and their post-binding fates differ. For example, while pIgR and Fcα/μR share D1, other domains differ. In mammals, Fcα/μR begins with a 46-residue N-terminal extension which was disordered in our structure. The Fcα/μR N-terminal extension is reminiscent of the 11-residue N-terminal extension found in avian pIgR, and structures containing *ap* SC revealed that a subset of those residues bridges interactions between the JC and SC domains^24,25^. Given that we do not observe ordered Fcα/μR N-terminal extension residues, it is likely regions preceding the first Ig-like domains in Fcα/μR and pIgR serve different roles. Another example is the pIgR C-terminal Ig-like domain (D4 in birds and D5 in mammals), which can form a covalent disulfide bond with both IgM and IgA. In contrast, we did not observe covalent interactions between mouse or mallard Fcα/μR and its ligands. This indicates that unlike pIgR, which delivers IgM or IgA to mucosal secretions and is cleaved, leaving SC bound to SIgM and SIgA, non-covalent high-affinity interactions between Fcα/μR and its ligands facilitate other functions and permit receptor recycling. Fcα/μR and pIgR also share a stalk region that connects ligand binding domains to the TM region. Both are predicted to be unstructured and anchor O-linked glycosylation; however, the *ap* pIgR stalk comprises 134 residues, approximately 11% of which have O-glycosylation potential whereas the ap Fcα/μR stalk spans 93 residues, 30% of which have O-glycosylation potential. Extensive O-linked glycosylation could promote a more rigid conformation, as demonstrated for clustered mucin domains^43^, and could protect Fcα/μR from proteolysis that typically cleaves the pIgR ectodomain from the cell membrane.

Cell type distribution of pIgR and Fcα/μR also differ; while pIgR is found on epithelial cells lining mucosal regions, Fcα/μR is found on a subset of hematopoietic cells (e.g. B-cells, macrophages) in lymphoid tissues and a subset of non-hematopoietic cells (e.g. follicular dendritic cells, mesangial cells). These geographic differences may eliminate any competition between pIgR and Fcα/μR binding to polymeric IgM and IgA ensuring different functional outcomes. Moreover, pIgR is thought to bind and transport largely free IgM and IgA, rather than antibody-antigen complexes. While as noted we observe robust interactions between Fcα/μR ectodomain and antigen-free IgA and IgM, it remains unknown how antigen binding impacts Fcα/μR’s down-stream functions.

## METHODS

### Identification of mallard *FCAMR*

To identify the mallard (Anas platyrhynchos) Fcα/μR homolog, the human Fcα/μR protein sequence (NCBI Accession: NP_001180214.1) was utilized as a query in a Basic Local Alignment Search Tool (tblastn)^48,49^ search. The search was restricted to the mallard genome using the Anas platyrhynchos organism filter (Taxonomy ID: 8839). This search identified a putative mallard *FCAMR* gene located on Chromosome 27 (NCBI Gene ID: 101802982), encoding the predicted protein sequence XP_005011823. Genomic location, structural organization, and annotation data were verified using the NCBI Gene database.

### RNA extraction and cDNA synthesis

All animal procedures were conducted in accordance with protocol 428, approved by the St. Jude Children’s Research Hospital Institutional Animal Care and Use Committee (IACUC). Five tissues (cloaca, bursa, esophagus, spleen, and small intestine) were collected from four mallard ducks (*Anas platyrhynchos*). Two of these birds had been previously infected with low pathogenicity avian influenza (LPAI) virus A/Mallard/Alberta/362/2017 (H3N8) at a dose of 10⁶ EID₅₀/mL and were euthanized on day 2 post-inoculation. Tissues were collected at necropsy and stored in 1 mL RNA protect (QIAGEN). Less than 30 mg of each tissue was homogenized using a Tissue Lyser II (QIAGEN), and total RNA was extracted using RNeasy kit (QIAGEN) according to the manufacturer’s instructions. Complementary DNA (cDNA) was synthesized from extracted RNA using the SuperScript IV First-Strand Synthesis System (Thermo Fisher Scientific) with random hexamers.

### PCR with gene-specific Primers

To ensure exclusive amplification of the target transcript, primers specific to mallard Fcα/μR were designed using NCBI Primer-BLAST^50^. To differentiate between amplified complementary DNA (cDNA) and potential genomic DNA (gDNA) contamination, PCR primers were specifically designed to span an intron–exon junction. Resulting PCR products were resolved using agarose gel electrophoresis, and the observed bands were excised and purified using Monarch^®^ DNA Gel Extraction Kit (New England BioLabs). Sequence identity of the purified amplicons was verified using Premium PCR service at Plasmidsaurus.

### Construct design

Protein sequences for the *Anas platyrhynchos* (*ap*; mallard duck) Fcα/μR (NCBI: XP_005011823.3), Mus musculus (*mm*; mouse) Fcα/μR (NCBI: NP_659209.2), *mm* IgM heavy chain constant region (HC) (Uniprot: P01872) were obtained. Target gene fragments were codon-optimized for human cells, synthesized (Integrated DNA Technologies), and cloned into the pD2610-v1 vector (Atum). The gene fragments generated were as follows: *ap* Fcα/μR ectodomain (18-436; NCBI: XP_005011823.3) was fused with exogenous Immunoglobulin heavy chain variable region VH26 signal peptide (MEFGLSWLFLVAILKGVQC; Uniprot: P01764) and a 6X Histag at C-terminus, *mm* Fcα/μR ectodomain (37-455; NCBI: NP_659209.2) was fused with exogenous tissue-type plasminogen activator signal peptide (MDAMKRGLCCVLLLCGAVFVSPAGA; Uniprot: P00750), followed by a thrombin cleavage site (LVPRGS), mCherry and C-terminal hexa-histidine tag. For *mm* IgM HC, Cμ2-Cμ3-Cμ4-Tp (105-454; Uniprot: P01872) was fused with endogenous signal peptide (MGWSWIFFFLLSGTAGVHC; translated from NCBI: BC096667.1) and a N-terminus 6X His tag. For *ap* IgM, IgA, JC and mm JC constructs as mentioned in our previous studies were used^24,25^. For SPR experiments, *ap* Fcα/μR were modified and synthesized to create following constructs: *ap* Fcα/μR^R225A-F229A^, *ap* Fcα/μR D1 (18-125; NCBI: XP_005011823.3) and *mm* Fcα/μR D1 (37-191; NCBI: NP_659209.2).

### Protein expression and purification

*ap* Fcα/μR and *mm* Fcα/μR DNA constructs were separately transfected into HEK Expi293 cells (Gibco: A14527) using the ExpiFectamine 293 Transfection Kit (Gibco: A14525). Four days after transfection, supernatants were harvested and the protein was purified by affinity chromatography using Ni-NTA Agarose (Qiagen) and eluted with elution buffer containing 50 mM Tris-HCl, 150 mM NaCl, pH 7.4 (1XTBS), and 250 mM imidazole. All proteins were further purified using size-exclusion chromatography (Superose 6 Increase 10/300 GL, Cytiva) and elution fractions containing protein were concentrated using Pierce™ Protein Concentrators (Thermo Fisher Scientific) and stored in 1XTBS at 4°C. *ap* FcμJ and *ap* FcαJ was expressed and purified as previously described^24,25^. *mm* FcμJ was purified using similar approach.

### Cryo-EM sample preparation and data collection

Protein complexes were prepared in 1× Tris-Buffered Saline (TBS) with continuous mixing by incubating individual purified proteins at room temperature for 1 hour followed by 24 hours at 4 °C. For the *ap* Fcα/μR–FcμJ complex, components were mixed at a 2:1 molar ratio. For the *ap* Fcα/μR– FcαJ and *mm* Fcα/μR–FcαJ complex, a 5:1 molar ratio was used. All three complexes were isolated via size-exclusion chromatography (Superose 6 Increase 10/300 GL, Cytiva) and verified by SDS-PAGE. The *ap* Fcα/μR–FcαJ complex was further purified using *ap* SC-coupled HiTrap NHS-Activated HP affinity column (Cytiva) to remove unbound antibody. All samples were pooled and concentrated using Pierce Protein Concentrators (Thermo Fisher Scientific).

Prior to sample vitrification, Quantifoil R1.2/1.3 Cu 400-mesh holey carbon grids (or UltraAuFoil R1.2/1.3 Au 300-mesh holey gold grids for ap Fcα/μR–FcαJ) were glow-discharged using a PELCO easiGlow system (Ted Pella Inc.) for 60 s at 25 mA. The *mm* Fcα/μR–FcμJ sample was bath-sonicated for 2 min immediately before grid application. Using a Vitrobot Mark IV (Thermo Fisher Scientific) operated at 4 °C and 100% relative humidity, 3 μL of sample was applied to the grids three consecutive times. Sample concentrations and blotting conditions were optimized as follows: 0.5 mg/mL with a 4.5 s blot time for *ap* Fcα/μR–FcμJ; 0.2 mg/mL with a 3 s blot time for *mm* Fcα/μR–FcμJ and *ap* Fcα/μR– FcαJ. All grids were blotted using a blot force of 2, and a 3 s wait time before plunge-freezing into liquid ethane.

Cryo-EM movies were recorded at Purdue University on a Titan Krios G4 transmission electron microscope (Thermo Fisher Scientific) operating at 300 kV, equipped with a post-GIF K3 direct electron detector (Gatan). Data acquisition was performed at 81,000× magnification in super-resolution mode using EPU software (Thermo Fisher Scientific) for untilted collections. For the *ap* Fcα/μR–FcμJ and *ap* Fcα/μR-FcαJ dataset, Leginon was additionally used for 30° stage-tilt collections. Total exposure times were 3.21 s, yielding a raw pixel size of 0.527 Å/pixel across 40 frames per movie. Total electron doses were 57.79 electrons/Å^2^ for *ap* Fcα/μR–FcμJ and 57.08 electrons/Å^2^ for *mm* Fcα/μR–FcμJ.

Data collection statistics can be found in Supplementary Table 2.

### Cryo-EM data processing, model building and refinement

All data processing was performed in cryoSPARC^51^ (v4.7 or higher). Imported movies (*ap* Fcα/μR– FcμJ: 4041 untilted, 2032 tilted; *mm* Fcα/μR–FcμJ: 4000 untilted; *ap* Fcα/μR–FcαJ: 12372 untilted, 7027 tilted) underwent motion correction, patch CTF estimation, and micrograph curation. Particles were picked using the blob picker (diameter: 150–300 Å, distance: 0.5) and extracted with a 2X Fourier crop. Curated particles were subjected to multiple rounds of 2D classification (four rounds for ap complexes; two rounds for the mm complex) before undergoing ab initio reconstruction and heterogeneous refinement.

Particles from the correct heterogeneous classes were re-extracted at the raw pixel size and subjected to non-uniform refinement^52^. For the *ap* Fcα/μR–FcαJ dataset, where density for the D2– D3 regions of the receptor degraded at higher contour levels, a global CTF refinement followed by a 3D classification was performed. The desired classes were pooled for homogeneous reconstruction, followed by non-uniform refinement. The final non-uniform refinements yielded maps with a global resolution of 3.27 Å for *ap* Fcα/μR–FcμJ, 3.9 Å for *ap* Fcα/μR–FcαJ and 4.13 Å for *mm* Fcα/μR–FcµJ at FSC = 0.143. Further local refinement of the *ap* Fcα/μR–FcαJ map yielded a resolution of 3.57 Å. The cryo-EM processing workflows for *ap* Fcα/μR–FcµJ, *ap* Fcα/μR-FcαJ and *mm* Fcα/μR–FcμJ are provided in Supplementary Figs. 2, 3, 4 and 5, respectively. Model validation results are provided in Supplementary Table S2.

Initial models of *ap* Fcα/μR, *mm* Fcα/μR and *mm* Cμ3-Cμ4 heavy chain domains were generated using the AlphaFold3 server. The generated models, *ap* FcμJ (PDB: 11DN) and ap FcαJ (PDB: 9EC7) were separately rigid body fitted into the refined map using ChimeraX^53^ v1.11.1, followed by iterative refinement in PHENIX^54^, COOT ^55,56^ v0.9.8.95, and ISOLDE^57^ v1.10.0. After individual refinements, the Fcα/μR and antibody ligand were combined and iteratively refined together. At the last round of refinement, glycans were built into the model, refined and validated using COOT and PHENIX software. Refinement statistics are shown in Supplementary Table 2.

### Sequence alignments

Multiple sequence alignments were generated using Clustal Omega^58,59^ v1.2.4 and visually rendered with ESPript 3.2^60^. Pairwise sequence identity and similarity percentages were calculated using EMBOSS Needle^61,62^ global pairwise alignment with default EBLOSUM62 matrix parameters.

### Structural analysis

Buried surface area at molecular interface was determined using PDBePISA^63^. Interchain contacts were determined using “contacts” tool of ChimeraX, having VdW overlap >= −0.4 Å, and PISA. To quantify the positional variations of Fcα/μR domains (D1–D3), structural alignment was done across three distinct pairs: (1) Fcα/μR–FcμJ versus SC–FcμJ, (2) Fcα/μR–FcμJ versus Fcα/μR–FcαJ, and (3) Fcα/μR–FcαJ versus SC–FcαJ. Superimpositions were performed in UCSF ChimeraX (version 1.11.1) utilizing two different alignment strategies depending on the structural conservation of the antibody backbone. For complexes sharing an identical antibody framework (comparisons 1 and 3), the Matchmaker tool was used to align the J chain. For comparisons across different antibody frameworks (comparison 2), structures were aligned using the D1 domain of Fcα/μR. Following alignment, global domain-specific displacements were calculated over Cα atoms using the “rmsd” command, which calculates the RMSD without iterative pruning so that the full magnitude of positional deviation relative to the reference state can be determined. For ap Fcα/μR Cα atoms used: D1 residues 1-108; D2 residues 118-218; D3 residues 229-325. For ap SC Cα atoms used: D1 residues: 13-120; D2 residues 130-230; D3 residues 241-337.

To compare domain topologies, structural alignments were performed using the Matchmaker tool in ChimeraX, focusing specifically on selected domain residues. Superimposition was refined through iterative pruning of atom pairs to calculate the RMSD of the structural cores. Interdomain orientations and rigid-body deviations were analyzed in ChimeraX. Principal structural axes for individual domains were generated via the “define axis” command using specified residues of domain boundaries. The angular variations of domains within a single receptor, or between homologous domains across superimposed complexes, with respect to each other were calculated using the “angle” command, which measures the geometric angle between the defined principal component vectors.

### Surface plasmon resonance

Surface plasmon resonance (SPR) binding studies were performed using a Sierra SPR-32 Pro (Bruker) operating at 25°C. Ligands (*ap* FcμJ, mm FcμJ, *ap* FcαJ, *ap* Fcμ and *ap* Fcα) were diluted in sodium acetate buffer pH 4.0 and immobilized on a High-Capacity Amine Sensor (Bruker) using an amine coupling kit (Bruker). Each ligand was immobilized in a separate flow channel with at least one mock-coupled spot per channel to serve as a reference surface.

A two-fold dilution series of each analyte was prepared using HBS EP+ buffer (10mM HEPES, 150 mM NaCl, 3mM EDTA, 0.05% (v/v) Tween 20) starting at 512 nM. Analytes were injected in HBS EP+ buffer at a flow rate of 50 μL/min, with a 120 s association phase and a 320 s dissociation phase. Each injection was followed by a regeneration step (2.5 M MgCl2, flow rate of 25 μL/min, 60 s contact time) and two wash steps (HBS EP+, 60 s contact time each).

Sensorgrams were reference-surface and buffer subtracted in Bruker Sierra Analyzer 3 software and then exported and replotted in Origin(Pro) version 2025. Normalization of sensorgrams for concentration-matched samples were carried out in Origin(Pro) by dividing each RU value by the maximum measured response of the respective analyte and then replotted in Origin(Pro). Sensorgrams shown in figures are representative of at least three replicate experiments.

To evaluate surface binding heterogeneity and kinetic profiles, raw data were fitted to a continuous surface-site distribution model using the EVILFIT software package^30^. The resulting kinetic rates and equilibrium dissociation constants were visualized as a 2D distribution heat map. Within this map, a boundary was defined around the target peak and integration of the signal-weighted amplitudes within this region yielded the final representative average K_D_.

### Figures

Structural figures were made using UCSF ChimeraX v1.11.1. To illustrate the binding footprints of Fcα/μR and SC on FcμJ or FcαJ, inter-chain contacts were identified using the ChimeraX “contacts” tool with a Van der Waals (VdW) overlap cutoff of >= −0.4 Å. The resulting interacting residues were mapped onto flat molecular surface representations of FcμJ or FcαJ, with the rest of the Fcα/μR or SC omitted to reveal only the surface of the receptor contacts. All graphs were plotted using Origin(Pro). All figures were assembled using Inkscape.

## ACKNOWLEDGEMENTS

The cryo-EM data collection was performed at the Purdue Cryo-EM Facility with the assistance of Thomas Klose and Frank Vago. Authors thank Profs. Wilfred van der Donk, Angad Mehta, Nicholas Wu, and Jenna Guthmiller, as well as Rebecca Schneider, Dalia Naser and all other members of the Stadtmueller lab for insightful discussions. Authors also thank Sarah Leonard for assistance with protein expression, DNA purification and assistance with SPR instrumentation.

## AUTHOR CONTRIBUTIONS

The study was conceived by BMS and MS; experiments and data analysis were conducted by MS; WH and RW provided the RNA samples from mallard tissues; EF assisted with DNA preparation and protein purification; and all authors contributed to manuscript writing.

## DATA AND MATERIALS AVAILABILITY

Cryo-EM density maps have been deposited in the EM databank (www.ebi.ac.uk/emdb) and the refined coordinate has been deposited in the Protein Data Bank (www.rcsb.org).

## Supplementary Materials

**Supplementary Fig. 1.**
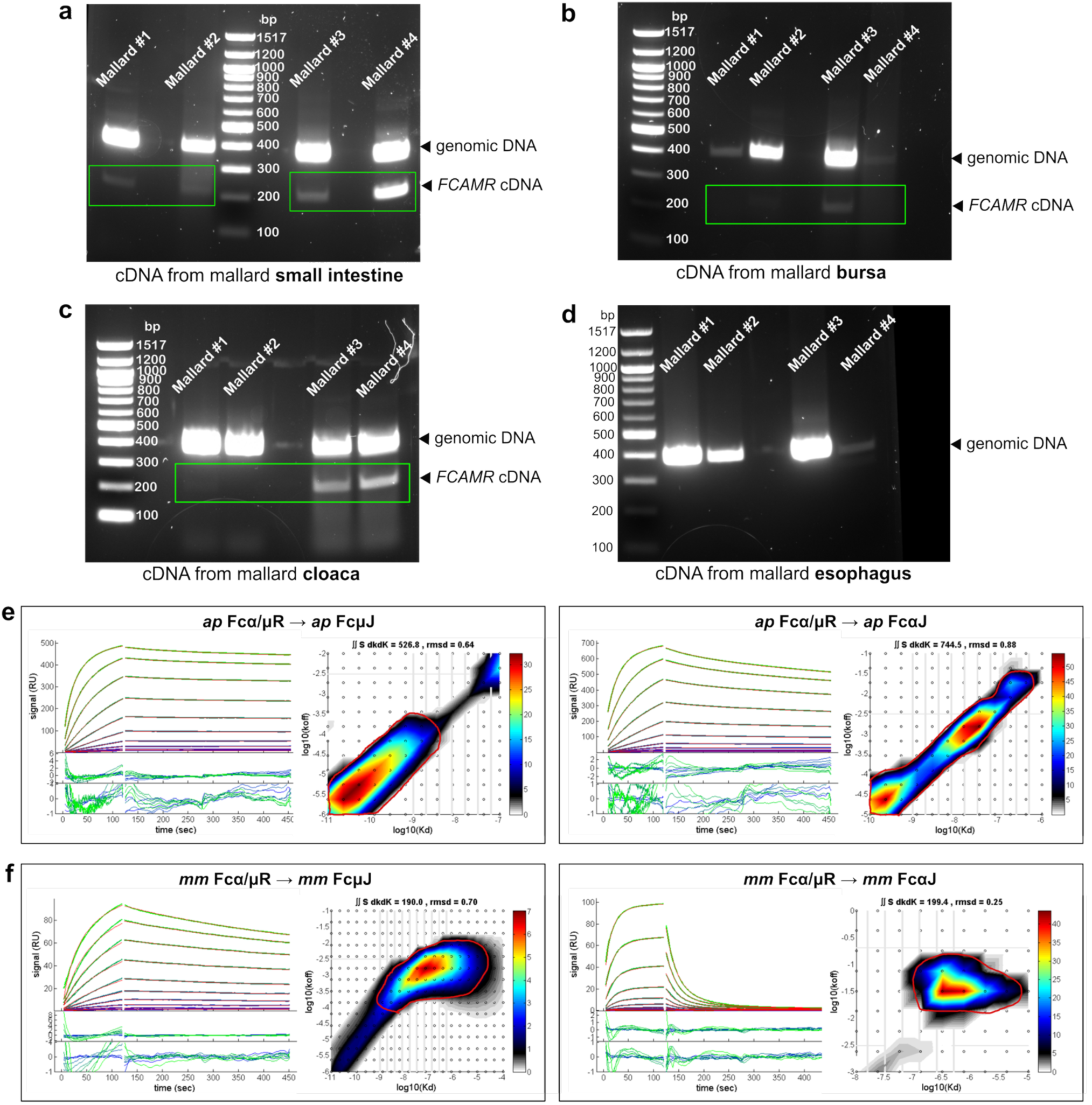
Tissue-specific expression of *ap* Fcα/μR, and *ap* and *mm* Fcα/μR interactions with FcμJ and FcαJ ligands. **(a-d)** 1.5% Agarose gel showing RT-PCR amplified *FCAMR* cDNA derived from the (**a**) small intestine, (**b**) bursa, (**c**) cloaca and (**d**) esophagus of four mallard ducks (Mallard #1–#4). The lower bands at 229 base pairs (bp) (green box) arise from amplified, spliced *FCAMR* mRNA. The upper bands arise from amplified unspliced genomic DNA. DNA ladders are indicated in bp. **(e-f)** SPR sensorgrams (left) and corresponding kinetic rate distributions (right) for the binding of (**e**) *ap* Fcα/μR to *ap* FcμJ (left box) and *ap* Fcα/μR to *ap* FcαJ (right box), and (**f**) *mm* Fcα/μR to *mm* FcμJ (left box) and *mm* Fcα/μR to *mm* FcαJ (right box). Experimental data (green lines) are overlaid with global kinetic fits (red lines) modeled using EVILFIT. Two-dimensional heatmaps (right panels) illustrate the continuous distribution of the log of affinity (K_D_) versus the dissociation rate constant (k_off_). The encircled region (in red) shows the area considered for calculating average K_D_.

**Supplementary Fig. 2.**
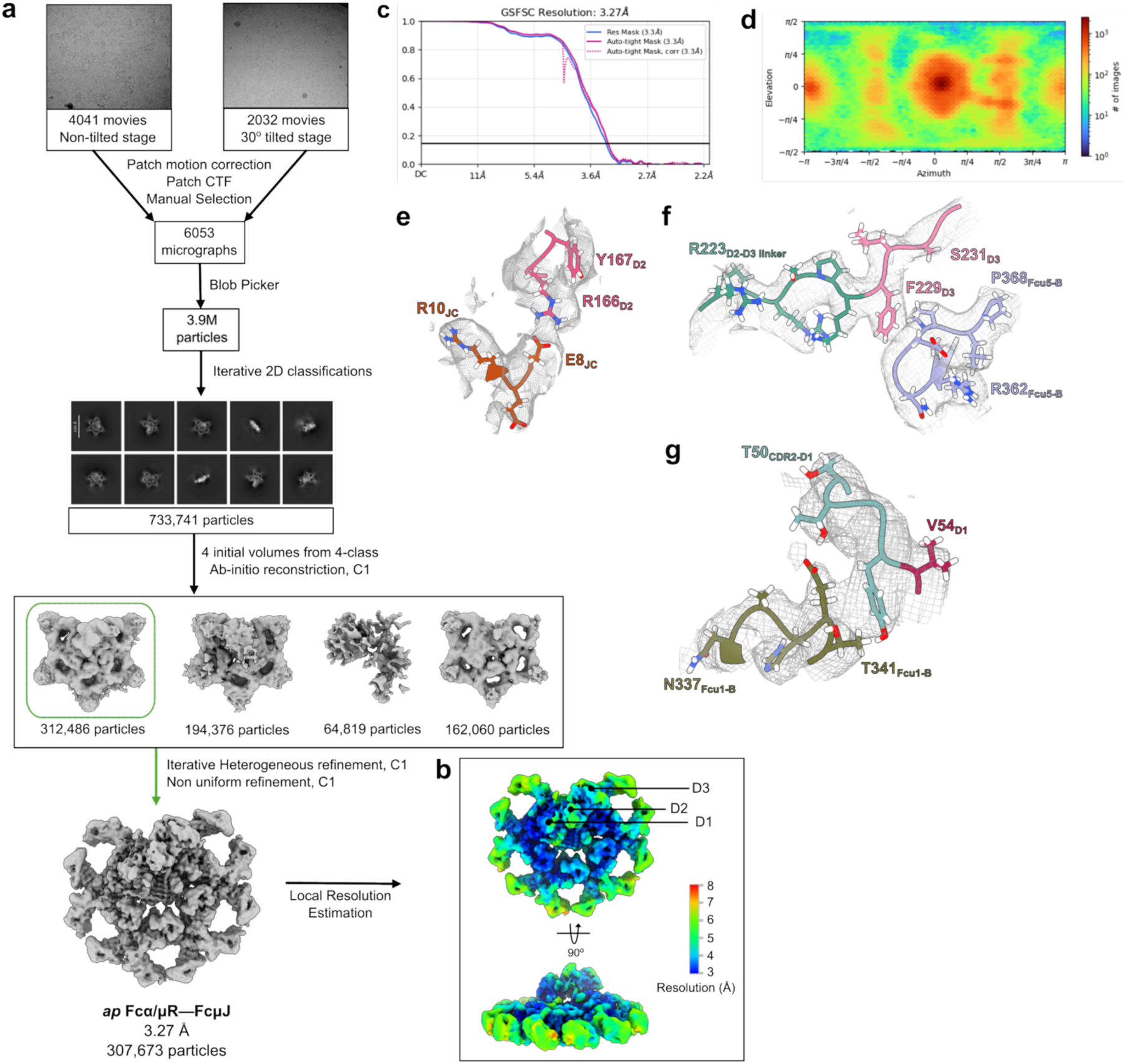
Cryo-EM workflow and structure determination of *ap* Fcα/μR–FcμJ complex. **(a)** Schematic summary of *ap* Fcα/μR—FcμJ data processing pipeline in CryoSPARC including representative micrographs of untilted and tilted data, and 2D class averages. **(b)** Local resolution map calculated by CryoSPARC and coloured in ChimeraX. **(c)** The FSC curve for the final reconstruction with reported resolution at FSC=0.143 shown by the horizontal line. **(d)** Viewing direction distribution plot of particles used in final refinement **(e)** Cryo-EM map density surrounding residues Fcα/μR D2 residues 166-167 and JC residues 8-10, contoured to threshold of 0.1 and carved at 2 Å. **(f)** Cryo-EM map density surrounding Fcα/μR D3 residues 223-231 and Fcμ5-B residues 362-368, contoured to threshold of 0.1 and carved at 2 Å. **(g)** Cryo-EM map density surrounding Fcα/μR D1 CDR2 residues 50-54 and Fcμ1-B residues 337-341, contoured to threshold of 0.1 and carved at 2 Å.

**Supplementary Fig. 3.**
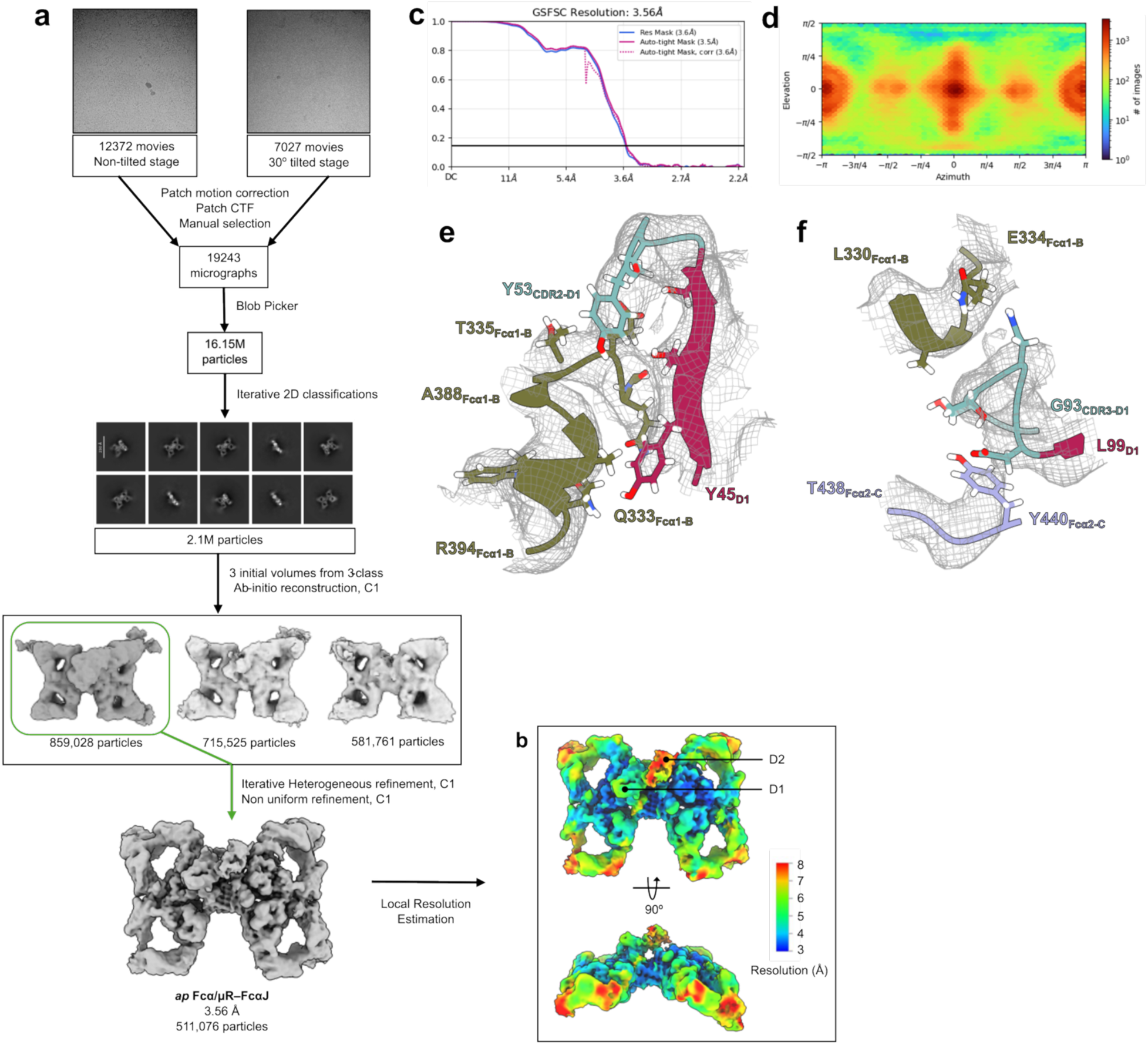
Cryo-EM workflow and structure determination of *ap* Fcα/μR–FcαJ complex. **(a)** Schematic summary of *ap* Fcα/μR–FcαJ data processing pipeline in CryoSPARC including representative micrographs of untilted and tilted data, and 2D class averages. **(b)** Local resolution map calculated by CryoSPARC and coloured in ChimeraX. **(c)** The FSC curve for the final reconstruction with reported resolution at FSC=0.143 shown by the horizontal line. **(d)** Viewing direction distribution plot of particles used in final refinement. **(e)** Cryo-EM map density surrounding *ap* Fcα/μR D1 CDR2 residues 45-53 and Fcα1-B residues 333-335, 388-394 contoured to threshold 0.22 and carved at 2Å. **(f)** Cryo-EM map density surrounding *ap* Fcα/μR D1 CDR3 residues 93-99, Fcα1-B residues 330-334, and Fcα2-C residues 438-440, each contoured to threshold 0.22 and carved at 2Å.

**Supplementary Fig. 4.**
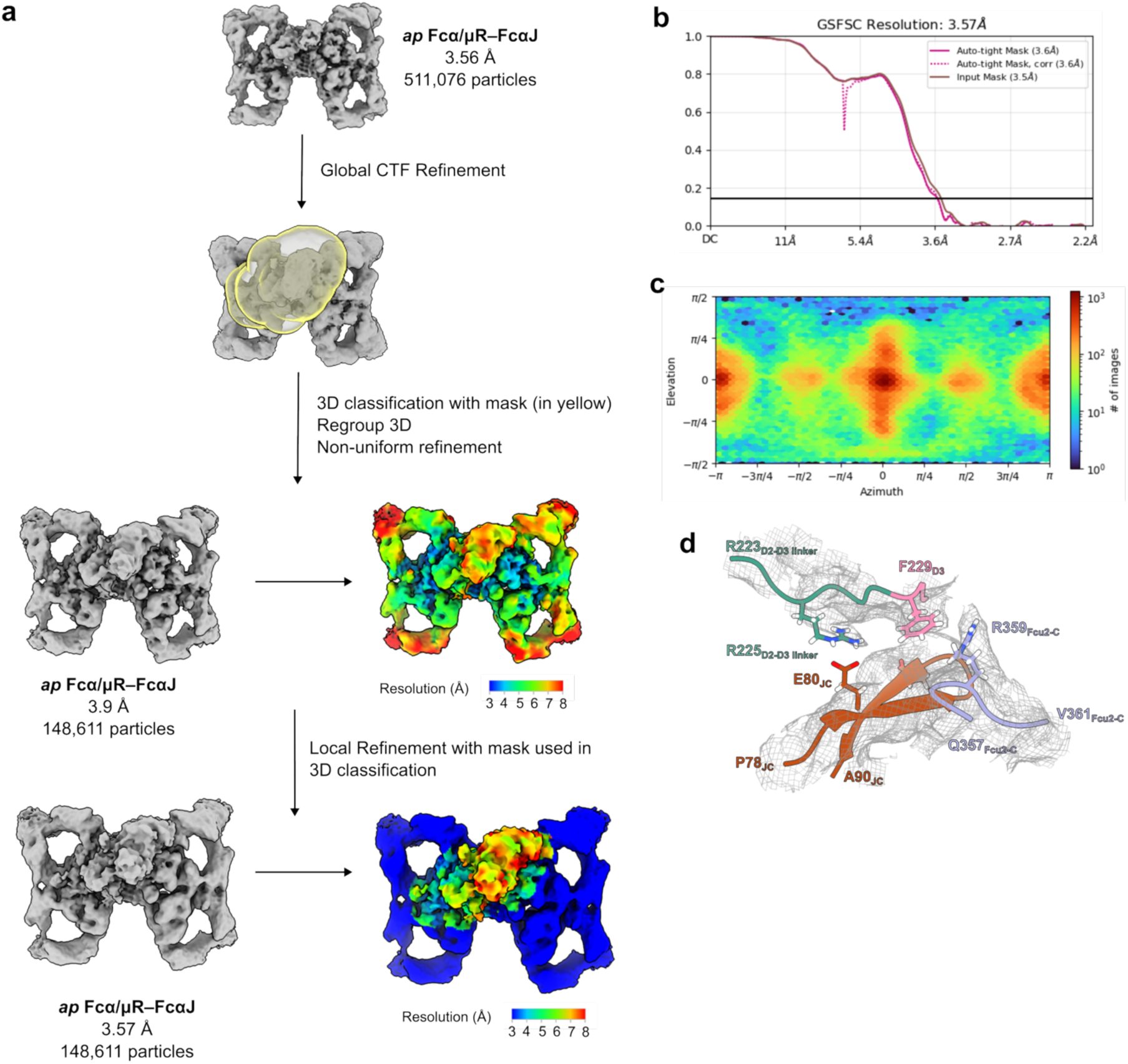
Focused Cryo-EM processing and map optimization of *ap* Fcα/μR D2–D3 region in complex with FcαJ. **(a)** Schematic summary of further *ap* Fcα/μR–FcαJ data processing pipeline in CryoSPARC to get a better density of D2-D3 region. The final map used is shown above while its local resolution map calculated by CryoSPARC and coloured in ChimeraX is shown below. **(b)** The FSC curve for the final reconstruction with reported resolution at FSC=0.143 shown by the horizontal line. **(c)** Viewing direction distribution plot of particles used in final refinement. **(d)** Cryo-EM map density surrounding *ap* Fcα/μR D3 residues 223-229, JC residues 78-90, and Fcα2-C residues 357-361, contoured to threshold 0.11 and carved at 2Å.

**Supplementary Fig. 5.**
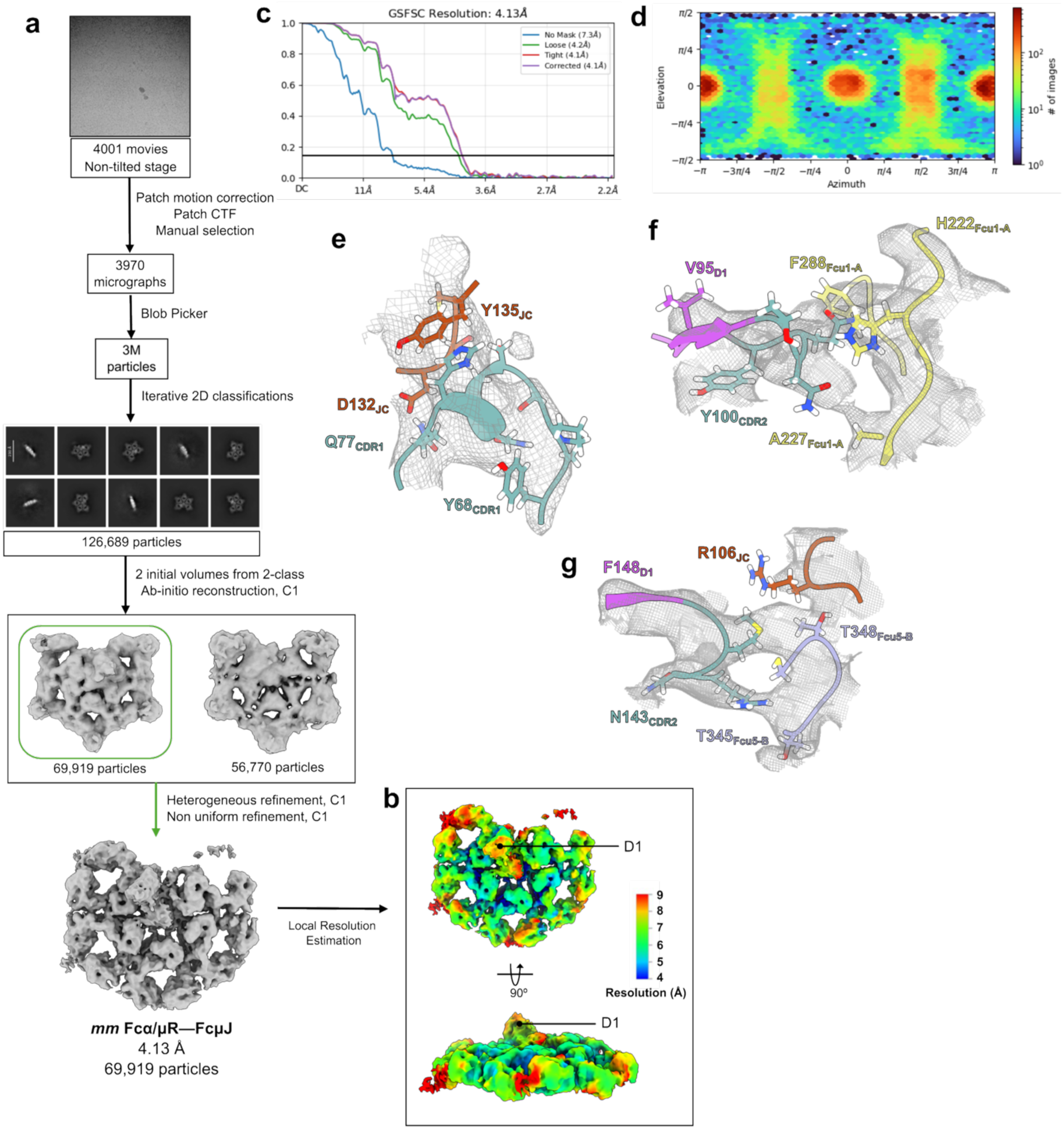
Cryo-EM workflow and structure determination of *mm* Fcα/μR–FcμJ complex. **(a)** Schematic summary of *mm* Fcα/μR–FcμJ data processing pipeline in CryoSPARC including representative micrographs of untilted and tilted data, and 2D class averages. **(b)** Local resolution map calculated by CryoSPARC and coloured in ChimeraX. **(c)** The FSC curve for the final reconstruction with reported resolution at FSC=0.143 shown by the horizontal line. **(d)** Viewing direction distribution plot of particles used in final refinement. **(e)** Cryo-EM map density surrounding *mm* Fcα/μR CDR1 residues 68-77 and JC residues 132-135, contoured to threshold 0.18 and carved at 3 Å. **(f)** Cryo-EM map density surrounding *mm* Fcα/μR CDR2 residues 95-100 and Fcμ1-A residues 222-227, 288, contoured to threshold 0.16 and carved at 2 Å. **(g)** Cryo-EM map density surrounding *mm* Fcα/μR CDR3 residues 143-148, JC residue 106 and Fcμ5-B residues 345-348, contoured to threshold 0.18 and carved at 2 Å.

**Supplementary Fig. 6.**
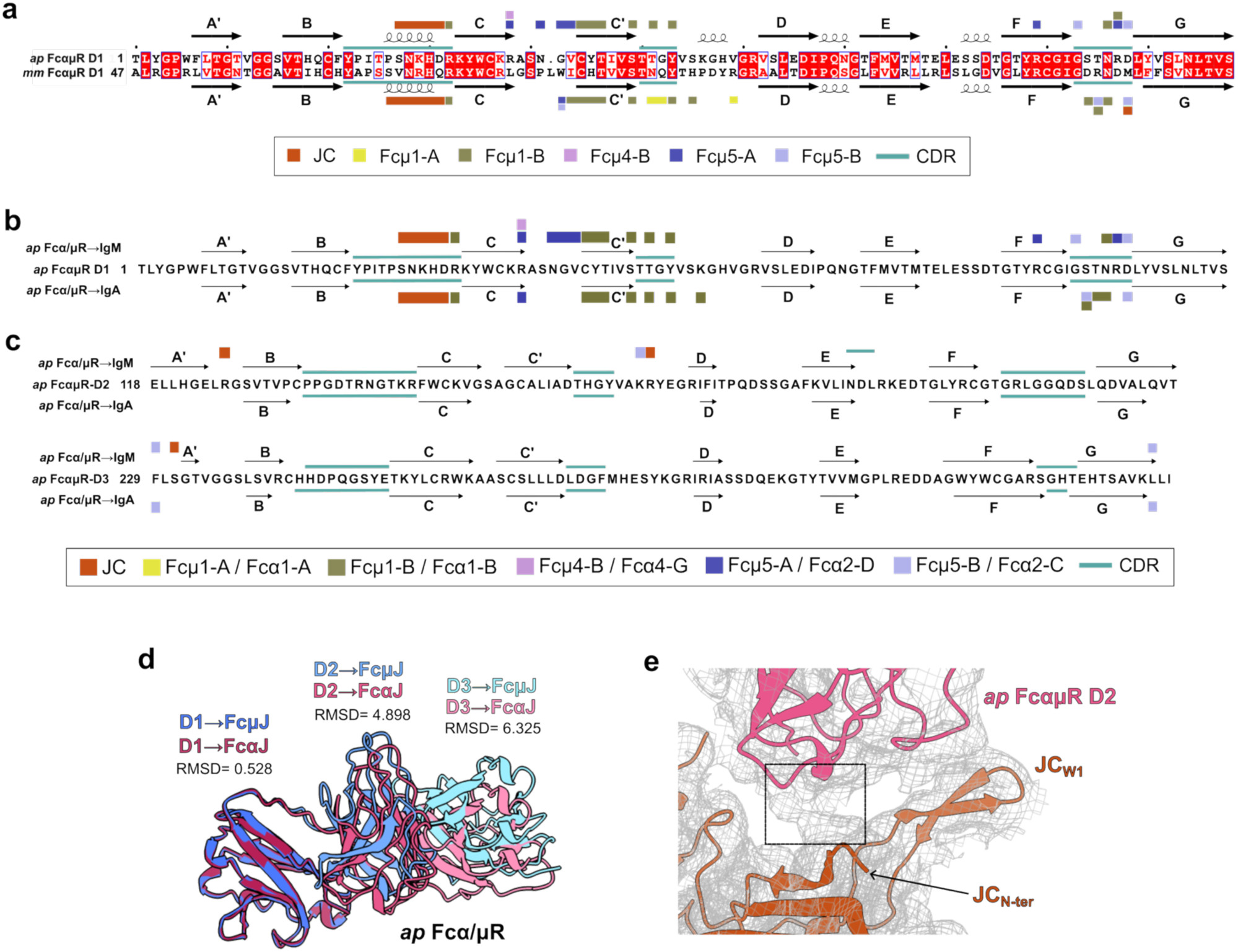
Sequence-based contact comparison of *ap* and *mm* Fcα/μR and structural analysis of *ap* Fcα/μR bound to IgM and IgA. **(a)** Sequence alignment of *ap* Fcα/μR D1 and *mm* Fcα/μR D1 highlighting residues that contact FcμJ (IgM). β-strands (A′–G) and CDR loops (cyan bars) are indicated. Colored boxes mark contacted target chains according to the key below. **(b)** *ap* Fcα/μR D1 sequence schematic highlighting residues that contact FcμJ (top) and FcαJ (bottom). β-strands (A′–G) and CDR loops (cyan bars) are shown, and colored boxes denote contacted target chains according to the key below. **(c)** *ap* Fcα/μR D2 and D3 sequence schematic highlighting residues that contact FcμJ (top) and FcαJ (bottom). β-strands (A′–G) and CDR loops (cyan bars) are shown, and colored boxes denote contacted target chains according to the key below. **(d)** Structural alignment of *ap* Fcα/µR bound to FcµJ (blue) versus FcαJ (pink). Individual domain alignments are labeled alongside their respective root-mean-square deviation (RMSD) values. **(e)** *ap* Fcα/μR–FcαJ cryo-EM density (grey mesh) and cartoon representation detailing potential *ap* Fcα/µR D2 contacts with the JC_N-ter_.

**Supplementary Fig. 7.**
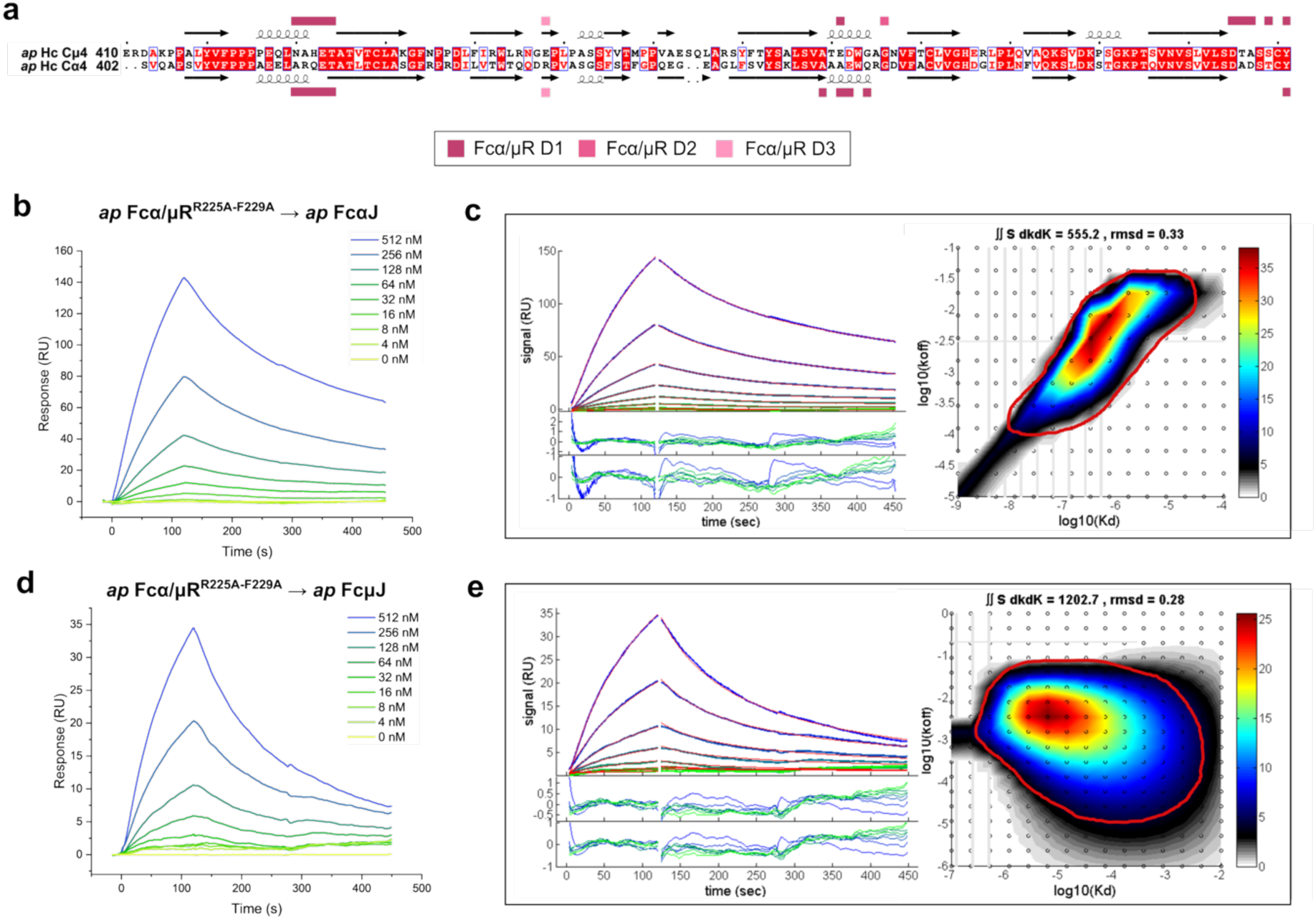
Contact residue mapping and binding analysis of the *ap* Fcα/μR mutant to IgM and IgA. **(a)** Sequence alignment of *ap* Cμ4 and Cα4 heavy chain constant domains, with conserved residues highlighted in red boxes. Colored squares positioned above and below the alignments indicate contact footprints of *ap* Fcα/μR domains (color-coded according to the key below). **(b-e)** SPR sensorgram (left) and corresponding fitted curves with kinetic rate distributions (in black box) for the binding of *ap* Fcα/µR^R225A-F229A^ to (**b,c**) *ap* FcαJ and to (**d,e**) *ap* FcµJ. Experimental data (green lines) are overlaid with global kinetic fits (red lines) modeled using EVILFIT, with fitting residuals shown on right (in black box). Two-dimensional heatmaps (right; black box) illustrate the continuous distribution of the log of affinity (K_D_) versus the dissociation rate constant (k_off_) calculated from the respective SPR sensorgrams. The encircled region (in red) shows the area considered for calculating average K_D_.

**Supplementary Fig. 8.**
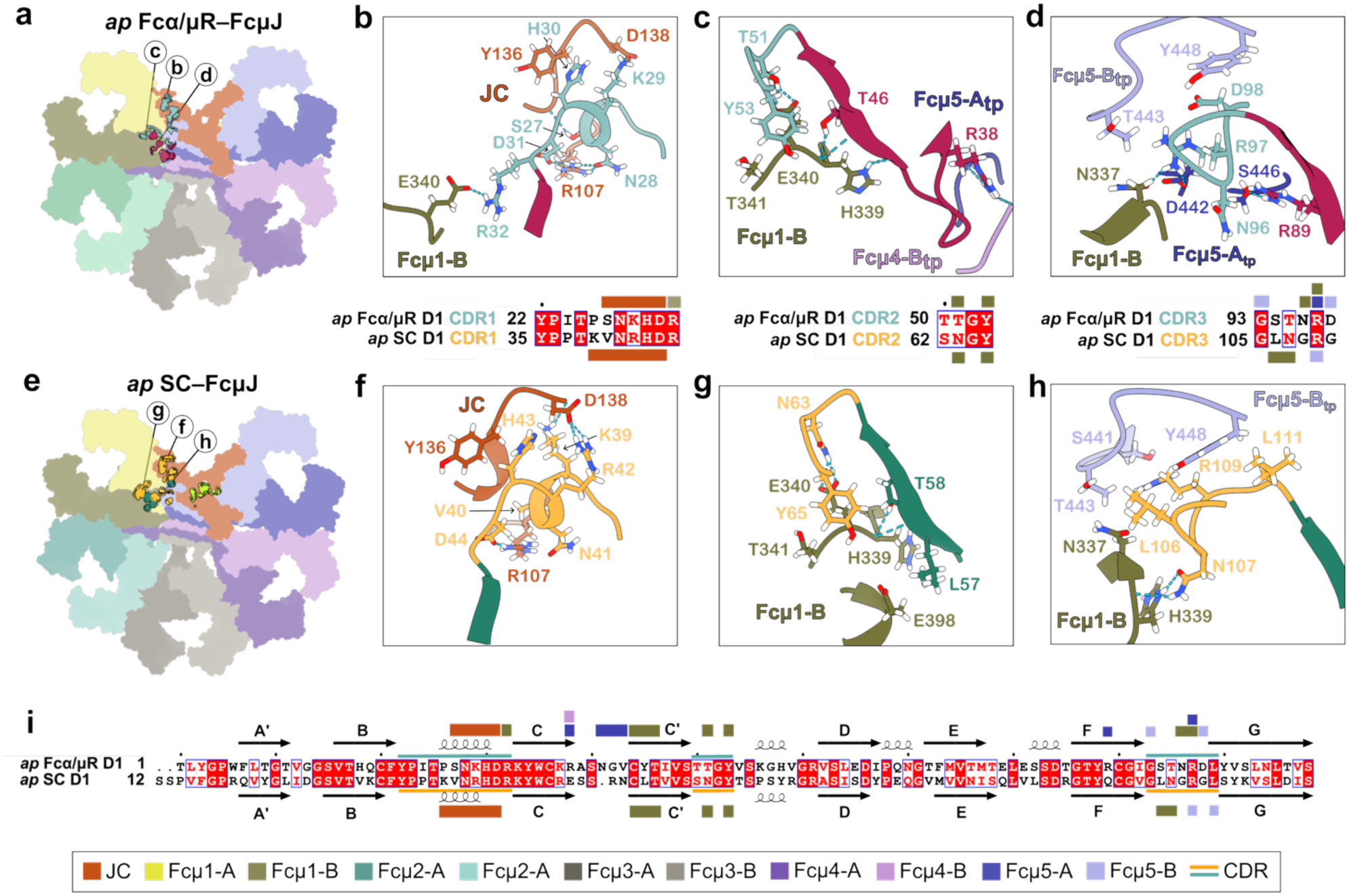
Comparative analysis of *ap* Fcα/µR D1 and *ap* SC D1 interactions with IgM. **(a)** *ap* FcµJ shown as a flat surface with the *ap* Fcα/µR D1 contact footprint shown in cyan (CDR residues) and raspberry (residues outside CDRs). Circled letters denote interfaces magnified in **b–d**. **(b–d)** (Top) Detailed views of *ap* Fcα/µR D1 CDR loops interacting with *ap* FcµJ at (**b**) CDR1, (**c**) CDR2, and (**d**) CDR3. Interacting residues are shown as sticks with hydrogen bonds and salt bridges indicated by dashed lines. (Bottom) Sequence alignments compare *ap* Fcα/µR D1 and *ap* SC D1 CDR loops, with contact footprints indicated by color-coded boxes (below Supplementary Fig. 8i). **(e)** *ap* FcµJ shown as a flat surface with the *ap* SC D1 contact footprint shown in yellow (CDR residues) and green (residues outside CDRs). Circled letters denote interfaces magnified in **f–h**. **(f–h)** Detailed views of *ap* SC D1 CDR loops interacting with *ap* FcµJ at (**f**) CDR1, (**g**) CDR2, and (**h**) CDR3. Interacting residues are shown as sticks with hydrogen bonds and salt bridges shown as dashed lines. **(i)** Structure-based sequence alignment comparing *ap* Fcα/µR D1 and *ap* SC D1 interaction with IgM. β-strands (A’–G) are indicated according to Ig fold nomenclature. Color-coded boxes indicate target chains contacted by specific residues (according to the key below).

**Supplementary Fig. 9.**
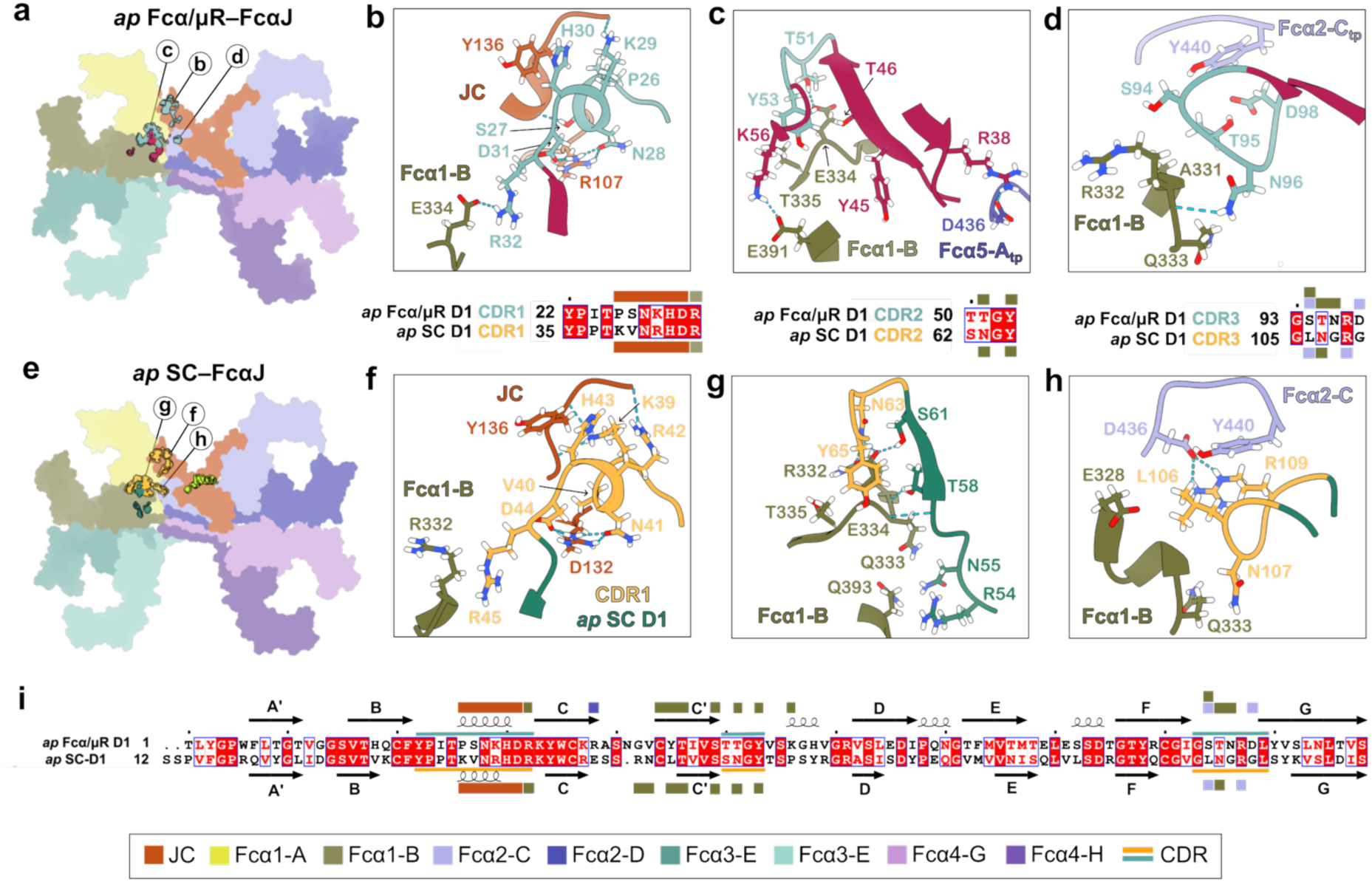
Comparative analysis of *ap* Fcα/µR D1 and *ap* SC D1 interactions with IgA. **(a)** *ap* FcαJ shown as a flat surface with the *ap* Fcα/µR D1 contact footprint shown in cyan (CDR residues) and raspberry (residues outside CDRs) Circled letters denote interfaces magnified in **b–d**. **(b–d)** (Top) Closeup views of *ap* Fcα/µR D1 CDR loops interacting with *ap* FcαJ at (**b**) CDR1, (**c**) CDR2, and (**d**) CDR3. Interacting residues are shown as sticks with hydrogen bonds and salt bridges indicated by dashed lines. (Bottom) Sequence alignments compare *ap* Fcα/µR D1 and *ap* SC D1 CDR loops, with contact footprints indicated by color-coded boxes (below Supplementary Fig. 9i). **(e)** *ap* FcαJ shown as a flat surface with the *ap* SC D1 contact footprint shown in yellow (CDR residues) and green (residues outside CDRs). Circled letters denote interfaces magnified in f–h. **(f–h)** Detailed views of *ap* SC D1 CDR loops interacting with *ap* FcαJ at (**f**) CDR1, (**g**) CDR2, and (**h**) CDR3. Interacting residues are rendered as sticks with hydrogen bonds and salt bridges shown as dashed lines. (**i**) Structure-based sequence alignment comparing *ap* Fcα/µR D1 and *ap* SC D1 interaction with IgA. β-strands (A’–G) are indicated according to Ig fold nomenclature. Color-coded boxes indicate target chains contacted by specific residues (according to the key below).

**Supplementary Fig. 10.**
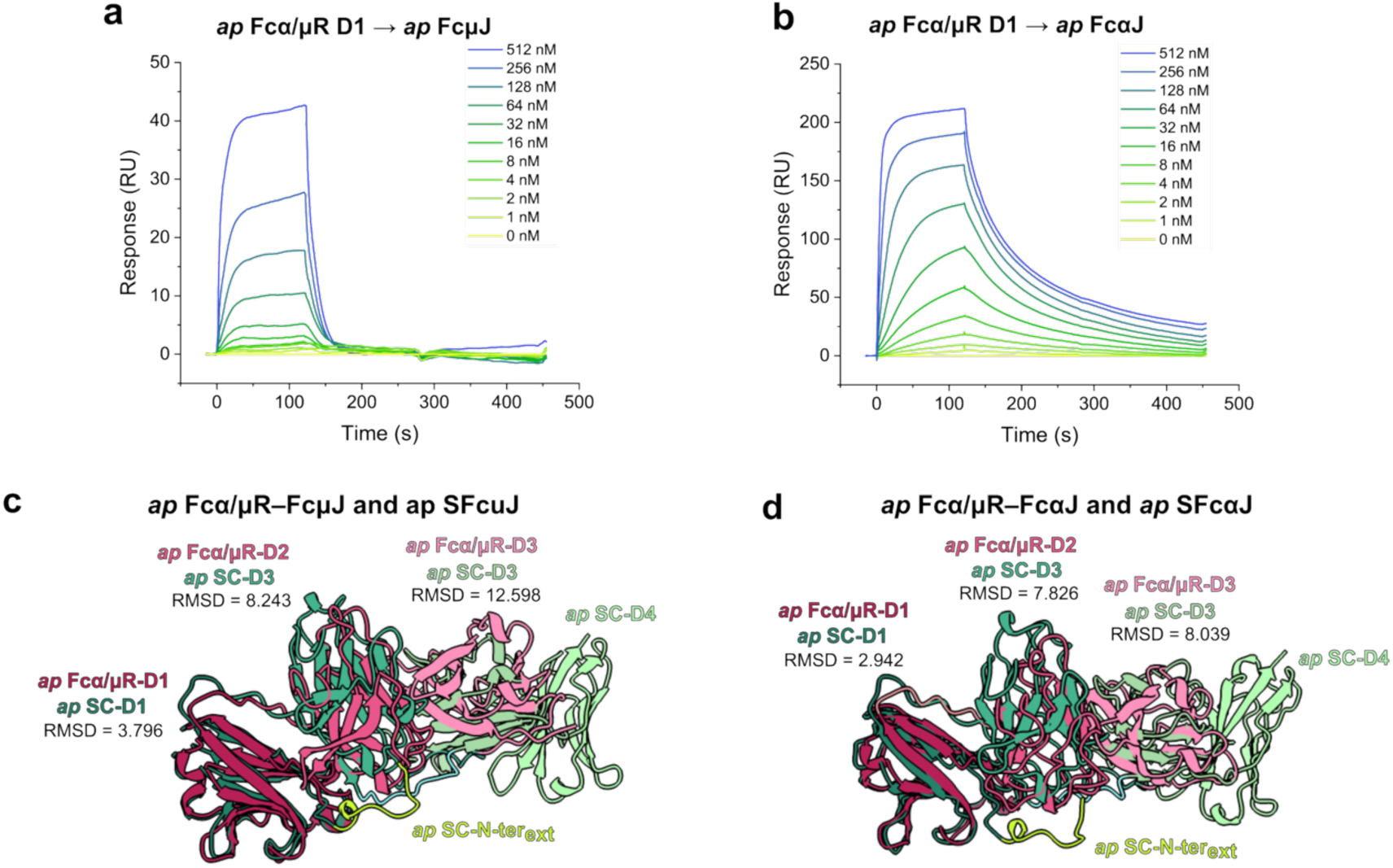
Binding affinity of *ap* Fcα/µR D1 and overall structural comparison of *ap* Fcα/µR and *ap* SC complexes. **(a, b)** SPR sensorgrams showing response units (RU) for a two-fold dilution series of: (**a**) *ap* Fcα/µR D1 binding to immobilized *ap* FcµJ, and (**b**) *ap* Fcα/µR D1 binding to immobilized *ap* FcαJ. (**c, d**) Structural superposition comparing *ap* Fcα/µR (pink) and *ap* SC (green) when bound to (**c**) FcµJ and (**d**) FcαJ, aligned on the JC. Domain-specific RMSD values between corresponding Ig-like domains are indicated.

**Supplementary Fig. 11.**
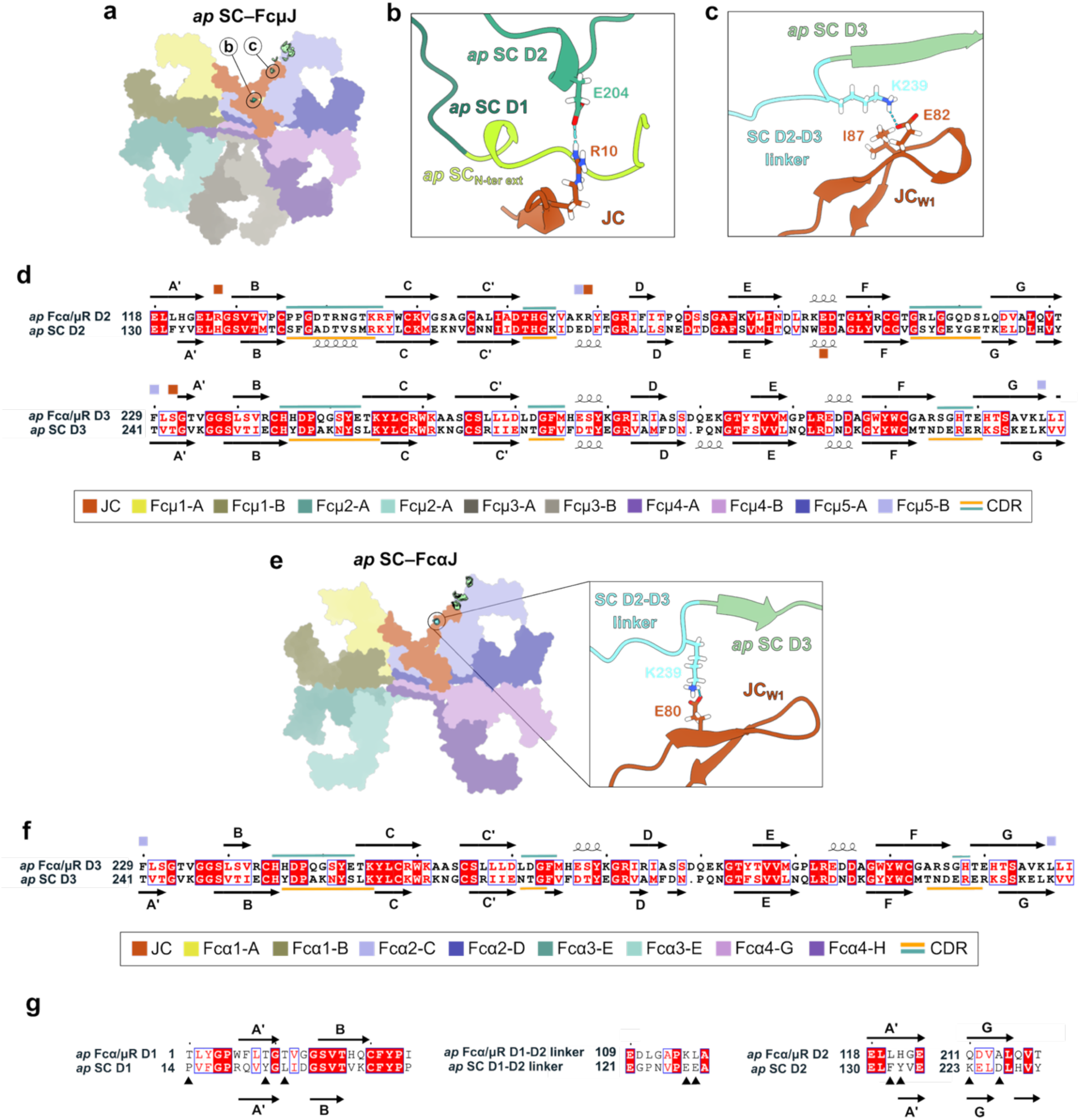
Comparison of *ap* Fcα/µR and *ap* SC D2–D3 interactions with IgM and IgA, alongside sequence divergence at the D1–D2 inter-domain interface. (**a**) *ap* FcµJ shown as a flat surface with the *ap* SC D3 (green) and D4 (light green) contact footprint shown and magnified in (b-c). (**b**) *ap* SC D2 and (**c**) *ap* SC D3 interactions with *ap* FcµJ. Key contact residues are shown as sticks with hydrogen bonds and salt bridges indicated by dashed lines. (**d**) Sequence alignment comparing *ap* Fcα/µR D2–D3 and *ap* SC D2–D3 interactions with IgM. β-strands are indicated (A’–G; according to Ig fold nomenclature). Color-coded boxes indicate target chains contacted by specific residues (refer to key). **(e)** *ap* FcαJ shown as a flat surface with the *ap* SC D3 (green) and D4 (light green) contact footprint shown. Magnified view of *ap* SC D2-D3 linker and D3 interactions with *ap* FcαJ is shown on right. **(f)** Sequence alignment comparing ap Fcα/µR D2–D3 and ap SC D2–D3 interactions with IgA. β-strands are indicated (A’–G; according to Ig fold nomenclature). Color-coded boxes indicate target chains contacted by specific residues (refer to key). **(g)** Sequence alignment of the D1–D2 interdomain interface and linker region between ap Fcα/μR and ap SC. Secondary structure elements (β -strands A′, B, and G) are indicated according to Ig-fold nomenclature. Black arrowheads highlight key residues that mediate the hydrophobic clamp and stabilizing electrostatic interactions at the D1–D2 interface in *ap* SC, which are replaced by smaller aliphatic, polar, or neutral residues in *ap* Fcα/μR.

**Supplementary Fig. 12.**
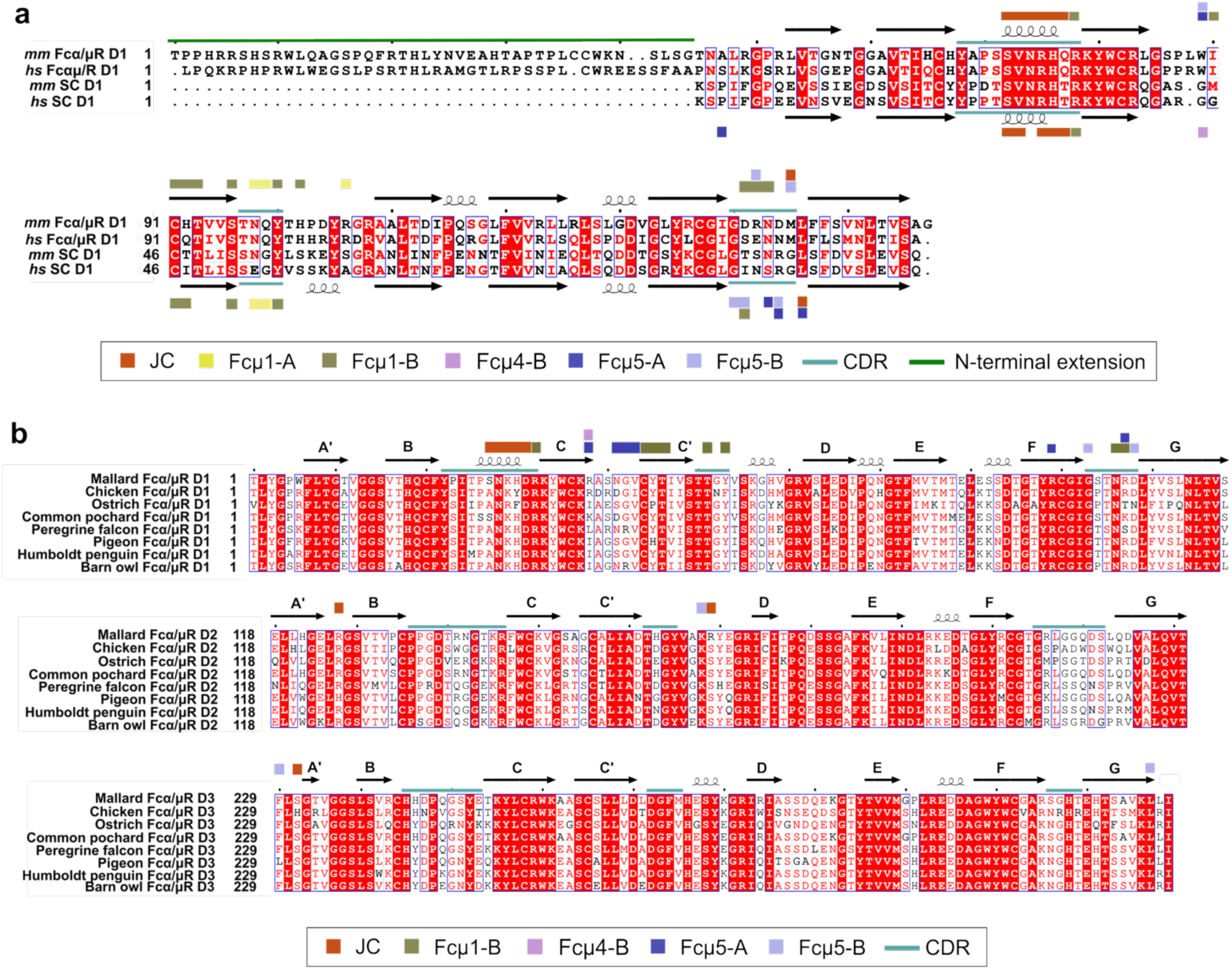
Sequence alignments comparing mammalian and avian Fcα/µR and SC contact residues. **(a)** Sequence alignments comparing D1 domains of mouse (*mm*) and human (*hs*) Fcα/µR and SC. Secondary structure elements (β-strands A’–G and loops) are indicated. Colored boxes above and below the sequences mark contact footprints corresponding to specific interacting chains according to the key below. **(b)** Sequence alignments comparing D1–D3 domains of avian Fcα/µR across representative species to indicate conservation of interacting Fcα/µR residues. Secondary structure elements (β-strands A’–G and loops) are indicated. Colored boxes above the mallard Fcα/µR sequence mark contact footprints of Fcα/µR with specific chains in FcµJ according to the key below.

**Supplementary Table 1.**
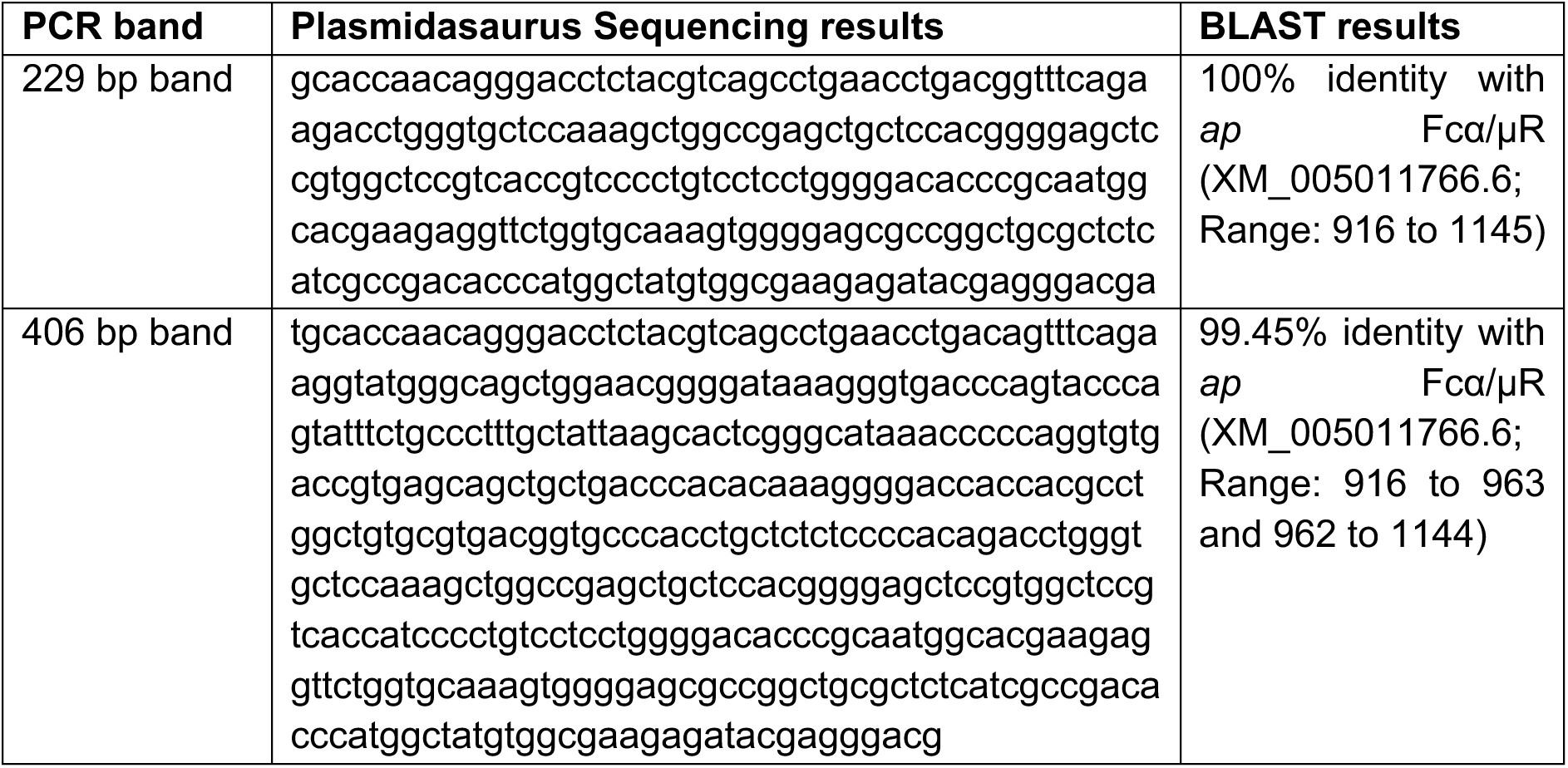
PCR sequence verification confirming ap Fcα/µR tissue expression.

**Supplementary Table 2.** Cryo-EM data collection and model refinement statistics.

|  | *ap* Fcα/μR–FcμJ | *mm* Fcα/μR–FcμJ | *ap* Fcα/μR–FcαJ |
|---|---|---|---|
| **Data collection** |  |  |  |
| **Voltage (kV)** | 300 | 300 | 300 |
| **Nominal magnification** | 81,000x | 81,000x | 81,000x |
| **Total dose (electrons/Å<sup>2</sup>)** | 57.79 | 57.08 | 57.79 |
| **Total exposure time (s)** | 3.21 | 3.21 | 3.21 |
| **Raw movie pixel size (Å/pixel)** | 0.527 | 0.527 | 0.527 |
| **Defocus range (μm)** | -0.6 – 1.8 | -0.6 – 1.8 | -0.6 – 1.8 |
| **Frames per movie** | 40 | 40 | 40 |
| **Automation software, 0°/30° tilt** | EPU/Leginon | EPU/-- | EPU/Leginon |
| **Data processing** |  |  |  |
| **Movies collected, 0°/30° tilt** | 4041 / 2032 | 4000 / 0 | 12372 / 7027 |
| **Initial particles** | 3.9M | 1.9M | 16.15M |
| **Final particles** | 307,673 | 69,919 | 148,611 |
| **Map resolution (Å)** | 3.27 | 4.13 | 3.57 |
| **FSC threshold** | 0.143 | 0.143 | 0.143 |
| **Model fit and validation** |  |  |  |
| **Model vs. data (CC)** |  |  |  |
| **Mask** | 0.82 | 0.77 | 0.79 |
| **Box** | 0.88 | 0.85 | 0.85 |
| **Peaks** | 0.78 | 0.70 | 0.70 |
| **Volume** | 0.81 | 0.77 | 0.78 |
| **Model composition** |  |  |  |
| **Total atoms** | 42632 | 36105 | 34057 |
| **Non-hydrogen atoms** | 21338 | 18143 | 17195 |
| **Protein residues** | 2744 | 2321 | 2246 |
| **Ligands** | BMA: 3; NAG:6 | — | BMA: 4; NAG: 9 |
| **Bond RMSD** |  |  |  |
| **Lengths (Å)** | 0.004 | 0.003 | 0.005 |
| **Angles (°)** | 0.601 | 0.599 | 0.605 |
| **MolProbity score** | 1.47 | 1.80 | 1.73 |
| **Clash score** | 4.48 | 6.24 | 6.09 |
| **Ramachandran plot (%)** |  |  |  |
| **Outliers** | 0.00 | 0.00 | 0.00 |
| **Allowed** | 3.68 | 5.01 | 5.89 |
| **Favored** | 96.32 | 94.99 | 94.20 |
| **Rotamer outliers (%)** | 0.67 | 1.40 | 0.79 |
| **Cβ outliers (%)** | 0.00 | 0.00 | 0.00 |
| **Cis proline** | 0.0 | 0.0 | 0.0 |
| **Twisted proline** | 0.0 | 0.0 | 0.0 |
| **CaBLAM outliers (%)** | 2.08 | 2.73 | 3.81 |

## REFERENCES

1. Flowers, E. M. et al. Identification of the Fc-alpha/mu Receptor in Xenopus provides insight into the emergence of the poly-Ig receptor (pIgR) and mucosal Ig transport. Eur. J. Immunol. 51, 2590 (2021).

2. Neves, F., de Sousa-Pereira, P., Melo-Ferreira, J., Esteves, P. J. & Pinheiro, A. Evolutionary analyses of polymeric immunoglobulin receptor (pIgR) in the mammals reveals an outstanding mutation rate in the lagomorphs. Front. Immunol. 13, 1009387 (2022).

3. Akula, S., Mohammadamin, S. & Hellman, L. Fc Receptors for Immunoglobulins and Their Appearance during Vertebrate Evolution. PLoS One 9, e96903 (2014).

4. Akula, S. & Hellman, L. The appearance and diversification of receptors for IgM during vertebrate evolution. Curr. Top. Microbiol. Immunol. 408, 1–23 (2017).

5. Shima, H. et al. Identification of TOSO/FAIM3 as an Fc receptor for IgM. Int. Immunol. 22, 149–156 (2010).

6. Wang, J. H. The sequence signature of an Ig-fold. Protein Cell 4, 569 (2013).

7. Zhang, R. et al. Xenopus IgX informs engineering strategies of IgM and IgG hexamers. Science Advances 11, 3737 (2025).

8. Hamburger, A. E., Bjorkman, P. J. & Herr, A. B. Structural insights into antibody-mediated mucosal immunity. Curr. Top. Microbiol. Immunol. 308, 173–204 (2006).

9. Johansen, Braathen & Brandtzaeg. Role of J chain in secretory immunoglobulin formation. Scand. J. Immunol. 52, 240–248 (2000).

10. Yoo, E. M., et al. Structural Requirements for Polymeric Immunoglobulin Assembly and Association with J Chain*. http://www.jbc.org (1999).

11. Lycke, N., Erlandsson, L., Ekman, L., Schön, K. & Leanderson, T. Lack of J Chain Inhibits the Transport of Gut IgA and Abrogates the Development of Intestinal Antitoxic Protection. The Journal of Immunology 163, 913–919 (1999).

12. Vaerman, J. P., Langendries, A., Giffroy, D., Brandtzaeg, P. & Kobayashi, K. Lack of SC/pIGR-mediated epithelial transport of a human polymeric IgA devoid of J chain: In vitro and in vivo studies. Immunology 95, 90–96 (1998).

13. Yoo, E. M., Trinh, K. R., Lim, H., Wims, L. A. & Morrison, S. L. Characterization of IgA and IgM binding and internalization by surface-expressed human Fcα/μ receptor. Mol. Immunol. 48, 1818–1826 (2011).

14. Shibuya, A., et al. Fcα/µ Receptor Mediates Endocytosis of IgM-Coated Microbes. http://immunol.nature.com (2000).

15. Sakamoto, N. et al. A novel Fc receptor for IgA and IgM is expressed on both hematopoietic and non-hematopoietic tissues. Eur. J. Immunol. 31, 1310–1316 (2001).

16. Kurita, N. et al. Identification of the Fcα/μR isoform specifically expressed in the kidney tubules. Mol. Immunol. 46, 749–753 (2009).

17. Arai, S. et al. Obesity-Associated Autoantibody Production Requires AIM to Retain the Immunoglobulin M Immune Complex on Follicular Dendritic Cells. Cell Rep. 3, 1187–1198 (2013).

18. Honda, S. I. et al. Marginal zone B cells exacerbate endotoxic shock via interleukin-6 secretion induced by Fcα/μR-coupled TLR4 signalling. Nat. Commun. 7, (2016).

19. Shibuya, A. & Honda, S. I. Molecular and functional characteristics of the Fcα/μR, a novel Fc receptor for IgM and IgA. Springer Seminars in Immunopathology vol. 28 377–382 Preprint at 10.1007/s00281-006-0050-3 (2006).

20. Kikuno, K. et al. Unusual biochemical features and follicular dendritic cell expression of human Fcα/μ receptor. Eur. J. Immunol. 37, 3540–3550 (2007).

21. Shibuya, A., Honda, S. I. & Shibuya, K. A pro-inflammatory role of Fcα/μR on marginal zone B cells in sepsis. International Immunology vol. 29 519–524 Preprint at 10.1093/intimm/dxx059 (2017).

22. Shibuya, A. & Honda, S.-I. Immune Regulation by Fca/l Receptor (CD351) on Marginal Zone B Cells and Follicular Dendritic Cells. (2015).

23. Schneider, R. M., Ji, Y. & Stadtmueller, B. M. Shared parts, different structures: How asymmetry drives IgM and IgA structure–function relationships. Curr. Opin. Struct. Biol. 101, 103346 (2026).

24. Schneider, R. M., Liu, Q. & Stadtmueller, B. M. The structures of tetrameric avian immunoglobulin A are uniquely structured to support mucosal immunity. bioRxiv 2025.09.19.677352 (2025) doi:10.1101/2025.09.19.677352.

25. Schneider, R. M., Liu, Q. & Stadtmueller, B. M. Cryo-EM Structure of Duck Secretory IgM Reveals a Conserved Pentameric Assembly with Avian-Specific Features at Molecular Interfaces. bioRxiv 2026.08.26.747385 (2026) doi:10.64898/2026.08.26.747385.

26. Stadtmueller, B. M. et al. The structure and dynamics of secretory component and its interactions with polymeric immunoglobulins. Elife 5, (2016).

27. Svitel, J., Boukari, H., Van Ryk, D., Willson, R. C. & Schuck, P. Probing the functional heterogeneity of surface binding sites by analysis of experimental binding traces and the effect of mass transport limitation. Biophys. J. 92, 1742–1758 (2007).

28. Svitel, J., Balbo, A., Mariuzza, R. A., Gonzales, N. R. & Schuck, P. Combined affinity and rate constant distributions of ligand populations from experimental surface binding kinetics and equilibria. Biophys. J. 84, 4062–4077 (2003).

29. Wyss, N. et al. Disentangling epitope spacing and antibody flexibility from avidity: A quantitative model for competition and bivalency in biosensor assays. Biophys. J. 125, 3214–3228 (2026).

30. Gorshkova, I. I., Svitel, J., Razjouyan, F. & Schuck, P. Bayesian Analysis of Heterogeneity in the Distribution of Binding Properties of Immobilized Surface Sites. Langmuir 24, 11577 (2008).

31. Chen, Q., Menon, R. P., Masino, L., Tolar, P. & Rosenthal, P. B. Structural basis for Fc receptor recognition of immunoglobulin M. Nature Structural & Molecular Biology 2023 30:7 30, 1033–1039 (2023).

32. Li, Y. et al. Immunoglobulin M perception by FcμR. Nature 2023 615:7954 615, 907–912 (2023).

33. Kumar, N., Arthur, C. P., Ciferri, C. & Matsumoto, M. L. Structure of the human secretory immunoglobulin M core. Structure 29, 564–571.e3 (2021).

34. Stadtmueller, B. M. et al. Biophysical and biochemical characterization of avian secretory component provides structural insights into the evolution of the polymeric Ig receptor. J. Immunol. 197, 1408 (2016).

35. Kumar Bharathkar, S. & Stadtmueller, B. M. Structural and Biochemical Requirements for Secretory Component Interactions with Dimeric IgA. The Journal of Immunology 213, 226–234 (2024).

36. Yang, X., Zhao, Q., Zhu, L. & Zhang, W. The three complementarity-determining region-like loops in the second extracellular domain of human Fc alpha/mu receptor contribute to its binding of IgA and IgM. Immunobiology 218, 798–809 (2013).

37. Bharathkar, S. K. et al. The structures of secretory and dimeric immunoglobulin a. Elife 9, 1–29 (2020).

38. Kaetzel, C. S. The polymeric immunoglobulin receptor: Bridging innate and adaptive immune responses at mucosal surfaces. Immunol. Rev. 206, 83–99 (2005).

39. Fulton, J. E. et al. The Chicken A and E Blood Systems Arise from Genetic Variation in and around the Regulators of Complement Activation Region. The Journal of Immunology 209, 1128–1137 (2022).

40. Oláh, I. & Glick, B. Avian lymph node: Light and electron microscopic study. Anat. Rec. 205, 287–299 (1983).

41. Lawn, A. M. & Rose, M. E. Presence of a complete endothelial barrier between lymph and lymphoid tissue in the lumbar lymph nodes of the duck (Anas platyrhynchos). Res. Vet. Sci. 30, 335–342 (1981).

42. McDonald, K. J., Cameron, A. J. M., Allen, J. M. & Jardine, A. G. Expression of Fc α/μ receptor by human mesangial cells: A candidate receptor for immune complex deposition in IgA nephropathy. Biochem. Biophys. Res. Commun. 290, 438–442 (2002).

43. Galashov, A. et al. Impact of Glycoclustering on Stiffening of MUC5AC Peptides Revealed by High-Efficiency Synthesis. Angew. Chem. Int. Ed Engl. 64, e202508278 (2025).

44. Takagaki, K., Satoh, K., Honda, S. I. & Shibuya, A. Molecular characterization of the dimer formation of Fcα/μ receptor (CD351). Mol. Immunol. 56, 23–27 (2013).

45. Kubagawa, H. et al. Physiological and Pathophysiological Roles of IgM Fc Receptor (FcµR) Isoforms. International Journal of Molecular Sciences 2023, Vol. 24, Page 5728 24, 5728 (2023).

46. Byland, R., Vance, P. J., Hoxie, J. A. & Marsh, M. A Conserved Dileucine Motif Mediates Clathrin and AP-2–dependent Endocytosis of the HIV-1 Envelope Protein. Mol. Biol. Cell 18, 414 (2007).

47. Kozik, P., Francis, R. W., Seaman, M. N. J. & Robinson, M. S. A Screen for Endocytic Motifs. Traffic 11, 843 (2010).

48. Gerts, E. M., Yu, Y. K., Agarwala, R., Schäffer, A. A. & Altschul, S. F. Composition-based statistics and translated nucleotide searches: Improving the TBLASTN module of BLAST. BMC Biology 2006 4:1 4, 41-(2006).

49. Altschul, S. F., Gish, W., Miller, W., Myers, E. W. & Lipman, D. J. Basic local alignment search tool. J. Mol. Biol. 215, 403–410 (1990).

50. Ye, J. et al. Primer-BLAST: a tool to design target-specific primers for polymerase chain reaction. BMC Bioinformatics 13, 134 (2012).

51. Punjani, A., Rubinstein, J. L., Fleet, D. J. & Brubaker, M. A. cryoSPARC: algorithms for rapid unsupervised cryo-EM structure determination. Nature Methods 2017 14:3 14, 290–296 (2017).

52. Punjani, A., Zhang, H. & Fleet, D. J. Non-uniform refinement: adaptive regularization improves single-particle cryo-EM reconstruction. Nature Methods 2020 17:12 17, 1214–1221 (2020).

53. Meng, E. C. et al. UCSF ChimeraX: Tools for structure building and analysis. Protein Science 32, e4792 (2023).

54. Liebschner, D. et al. Macromolecular structure determination using X-rays, neutrons and electrons: recent developments in Phenix. urn:issn:2059-7983 75, 861–877 (2019).

55. Emsley, P., Lohkamp, B., Scott, W. G. & Cowtan, K. Biological Crystallography Features and development of Coot. 10.1107/S0907444910007493 doi:10.1107/S0907444910007493.

56. Emsley, P. & Cowtan, K. Coot: model-building tools for molecular graphics. urn:issn:0907-4449 60, 2126–2132 (2004).

57. Croll, T. I. ISOLDE: a physically realistic environment for model building into low-resolution electron-density maps. urn:issn:2059-7983 74, 519–530 (2018).

58. Sievers, F. et al. Fast, scalable generation of high-quality protein multiple sequence alignments using Clustal Omega. Mol. Syst. Biol. 7, (2011).

59. Madeira, F. et al. The EMBL-EBI Job Dispatcher sequence analysis tools framework in 2024. Nucleic Acids Res. 52, W521 (2024).

60. Gouet, P., Courcelle, E., Stuart, D. I. & Métoz, F. ESPript: analysis of multiple sequence alignments in PostScript. Bioinformatics 15, 305–308 (1999).

61. Rice, P., Longden, L. & Bleasby, A. EMBOSS: The European Molecular Biology Open Software Suite. Trends in Genetics 16, 276–277 (2000).

62. Needleman, S. B. & Wunsch, C. D. A general method applicable to the search for similarities in the amino acid sequence of two proteins. J. Mol. Biol. 48, 443–453 (1970).

63. Krissinel, E. & Henrick, K. Inference of Macromolecular Assemblies from Crystalline State. J. Mol. Biol. 372, 774–797 (2007).

